# Chemical augmentation of the validated HepaRG^TM^ CYP enzyme induction test method Part 1: The Goliath two laboratory study

**DOI:** 10.64898/2026.06.16.732540

**Authors:** Miriam N Jacobs, Barbara Kubickova, Elodie Person, Jorke H Kamstra, Nicolas Cabaton, Sebastian Hoffmann, Agnès Jamin, Marlene Lacroix, Juliette Legler, Vesna Munic Kos, Sandra M Nijmeijer, Theo L Sinnige, Laurie Urien, Daniel Zalko

## Abstract

Cytochrome P450 (CYP) enzymes play a key role in the metabolism of both xenobiotics and endogenous compounds, and the activity of some CYP isoforms are susceptible to induction and/or inhibition by certain chemicals. As CYP induction and inhibition can significantly alter the *in vivo* fate of xenobiotics i.e., levels of parent chemicals and/or metabolites, and thus toxicity, CYP induction/inhibition data is needed for regulatory chemical toxicity hazard assessment.

Utilizing available human *in vivo* pharmaceutical data, a successful validation was previously conducted on the *in vitro* HepaRG™ CYP induction test method for measurement of induction of three key human CYP enzymes CYP1A1/1A2, 2B6 and 3A4. However, further validation data was required to demonstrate applicability of the test method to also accurately detect CYP induction mediated by industrial and pesticidal chemicals. Here we report on the supplementary validation of the HepaRG™ CYP enzyme induction test method carried out in two laboratories under the auspices of the EU Horizon2020-funded project “GOLIATH”, to expand the chemical applicability domain beyond pharmaceutical chemicals. Successful transfer was demonstrated and reproducibility assessed for the original 10 selected proficiency pharmaceuticals, plus three reference inducers together with six additional non-pharmaceutical ‘augmentation chemicals’. The method and chemical selection were found to be reliable and relevant for the routine assessment of human CYP induction. For the augmentation chemicals being proposed as additional proficiency chemicals, the test method achieved a reasonable but not optimum reproducibility. Recommendations are proposed to improve the test method’s specificity, reflecting the inherent uncertainty around borderline CYP inducing chemicals.

**Plain language summary:** Cytochrome P450 (CYP) enzymes help break down drugs and other chemicals in the body. Their activity can be increased (induced) or decreased (inhibited), which can change how toxic a chemical is and when it is excreted. Because of this, CYP data is important for chemical safety assessments. A laboratory-based method using HepaRG cells was previously validated to measure induction of key CYP enzymes (CYP1A1/1A2, CYP2B6 and CYP3A4) using pharmaceutical chemicals. This study aimed to show that it also works well for industrial and pesticidal chemicals. In the EU funded GOLIATH project, two laboratories tested 10 pharmaceutical and 6 non-pharmaceutical chemicals. The method showed good reliability overall and strong reproducibility for pharmaceuticals. For non-pharmaceutical chemicals, results were acceptable but less consistent. The study concludes that the method is useful for routine testing, but improvements are needed to increase accuracy and better handle chemicals that show weak or borderline CYP induction effects.

## 1 Introduction and background

Cytochrome P450 (CYP) enzyme induction represents a sensitive biomarker for phenotypic metabolic competence of *in vitro* systems. Due to their abundance, inducibility and functional versatility, CYP enzymes are particularly important in xenobiotic metabolism. Substrates can either be catalytically detoxified or bioactivated. CYP enzyme induction, covering both *de novo* protein synthesis and protein stabilization, is the functional endpoint of CYP induction and the basis of potential chemical interactions in humans (Lewis, 2002). CYP enzyme induction is also a molecular initiating event (MIE) and Key Event (KE) in disease and a biomarker for the activation of toxicologically relevant nuclear receptor signalling pathways, including the Aryl hydrocarbon Receptor (AhR; CYP1A1/1A2), the Pregnane X Receptor (PXR; CYP3A4), and the Constitutive Androstane Receptor (CAR; CYP2B6). Furthermore, alterations to the activity levels of CYP enzymes can have a direct impact upon the level and chemical nature of exposure to xenobiotics and metabolites. Similarly, when considered on the basis of their endogenous substrates (such as hormones), this may cause dysregulation of normal metabolism and homeostasis, including energy related metabolism, with potential adverse effects.

CYP induction can be both modelled and measured in animal experiments *in vivo*. However, it is well documented that the induction/inhibition profiles and in AhR, PXR and CAR activation can differ substantially between species such as between rodents and humans for instance, with major toxicokinetic implications.

The scientific basis and regulatory need for a human *in vitro* CYP (enzyme) induction test method has been well documented in the scientific literature and by international organizations for a long time now (e.g. Coecke et al. 2006, Jacobs et al. 2008 OECD 2008). The *in vitro* HepaRG CYP induction test method was identified as the most promising method to take forward for validation for OECD Test Guideline (TG) purposes (OECD 2008, Jacobs et al. 2013), and the European Commission’s Joint Research Centre (JRC) Reference Laboratory (EURL) European Centre for the Validation of Alternative Methods (ECVAM) duly embarked upon a validation study to address this need (Bernasconi et al. 2019).

The *in vitro* CYP enzyme induction interlaboratory validation study (Bernasconi et al. 2019, JRC 2014a) compared the performance of cryopreserved primary human hepatocytes (PHHs) and differentiated human HepaRG™ cells. Whilst the CYP enzyme activity and gene expression profile of PHHs might be more representative of the human *in vivo* situation, the inherent higher biological variability of PHH preparations, as demonstrated in the validation study (JRC, 2014a), and assessed in the subsequent peer review (JRC 2014b, 2014c), reduces reproducibility and consequently regulatory utility. The cryopreserved differentiated HepaRG™ cells were therefore considered more appropriate for OECD Test Guideline (TG) development.

With respect to the choice of CYP isoforms that are evaluated in this test method, the four P450 iso-enzymes 1A2, 2B6, 2C9 and 3A4 were initially selected for the EURL ECVAM’s validation study, as they are primary CYPs that are inducible in humans. These isoforms are involved in many of the Phase I detoxifying processes in human liver and are recommended by the European Medicines Agency (EMA) and the US Food and drug Administration (FDA) drug-drug interaction Guidelines (EMA 2012; 2024, US FDA 2020 and 2024). CYP3A4 is widely considered to be one of the most abundant isoforms, constituting 30 % of all the CYP liver enzymes in humans. The CYP2C family accounts for 30-40 % of human hepatic CYPs, with CYP2C9 being the most highly expressed. However, during the EURL ECVAM validation study, CYP2C9 inducibility did not fulfil the acceptance criteria with the inducer positive control (10 µM rifampicin) and frequently showed flat/variable concentration-response curves. For this reason, measurement of CYP2C9 activity was excluded from the test method (JRC 2014c). This is also in line with both CYP3A4 and CYP2C9 being induced via activation of PXR and clinical evidence on CYP2C9 induction (by rifampicin) being much lower than CYP3A4 induction, and consequently the US FDA (FDA 2024) and EMA (EMA 2012) also do not include CYP2C9 in the CYP induction test battery. EMA only requires the potential for CYP2C8, CYP2C9, and CYP2C19 induction to be investigated if the test pharmaceutical “*has been shown to be an inducer of CYP3A4 in a clinical study*” (EMA 2024).

The HepaRG™ CYP induction test method standard operating protocol (SOP) was first provided in 2009 (JRC Test Method Number: TM2009-14 (EU), Protocol/SOP N°194, and was validated by EURL-ECVAM using pharmaceuticals (JRC 2014 a,b, c, Bernasconi et al. 2019). The test method investigates the potential of chemicals to induce the catalytic activity of key CYP enzymes (CYP1A2, CYP3A4 and CYP2B6, mediated via AhR, PXR or CAR) in HPR116 cells (differentiated HepaRG cells). It consists of a 48-h exposure to a test chemical, appropriate positive controls and solvent/vehicle controls (with three technical replicates), followed by the addition of a cocktail of CYP isoform specific probe substrates for the three target CYP enzyme isoforms and subsequent liquid chromatography/mass spectrometry (LC/MS) analyses and quantification of probe metabolites accounting for key CYP enzyme activities. Following the analyses of the results, after conducting the test method on at least three different cell batches (biological replicates) it is then possible to determine if a chemical is an inducer for a specific CYP enzyme, or not.

However, in 2019 this validation study (JRC 2014, Bernasconi et al. 2019), the peer review report (JRC 2014b, 2014c) and the draft TG (no longer available) were not accepted by the OECD Working Group of National Coordinators to the OECD Test Guideline Programme (WNT), as the need for the inclusion of a broader chemical applicability domain in the proficiency chemicals, was expressed, including non-pharmaceutical chemicals, despite a lack of suitable and available human *in vivo* data. To take these needs into account, a different methodology was needed to be able to develop an extension to the chemical selection, with which to augment the applicability domain. This proposed methodology was first formally agreed with the WNT in April 2021 (also on the basis of prior WNT discussions) and then chemical selection work was undertaken based upon a selected review of the scientific literature (Jacobs et al. 2022a).

Selection of the candidate non-pharmaceutical proficiency chemicals (“augmentation chemicals”) was made in accordance with accepted guidance for the validation of screening-type test systems (Hartung et al. 2004; OECD 2005; OECD 2018). A long list of 23 non-pharmaceutical chemicals was first identified as additional potential candidate proficiency chemicals, on the basis of specific CYP induction profiles and availability of data from humans/human based test systems as previously reported (Jacobs et al. 2022a). From this, a short list of 6 augmentation chemicals was selected (Tab. 1, Augmentation chemicals). Preliminary chemical potency classification for the augmentation chemicals was based upon the induction of CYP enzyme catalytic activity data. Gene expression induction was not considered to be of sufficient strength to allow provisional potency classification (discriminating “no”, “low”, “moderate” and “strong” inducers) and was used as supporting evidence only. Similarly, evidence from human biomonitoring studies was included as supporting (not primary) evidence for human *in vivo* metabolism due to e.g., a lack of defined exposure scenarios in such studies. Suitable negative (non-inducer) chemicals were identified, to allow evaluation of the specificity of a test system, in accordance with test method validation guidance (OECD 2005).

**Tab. 1.**
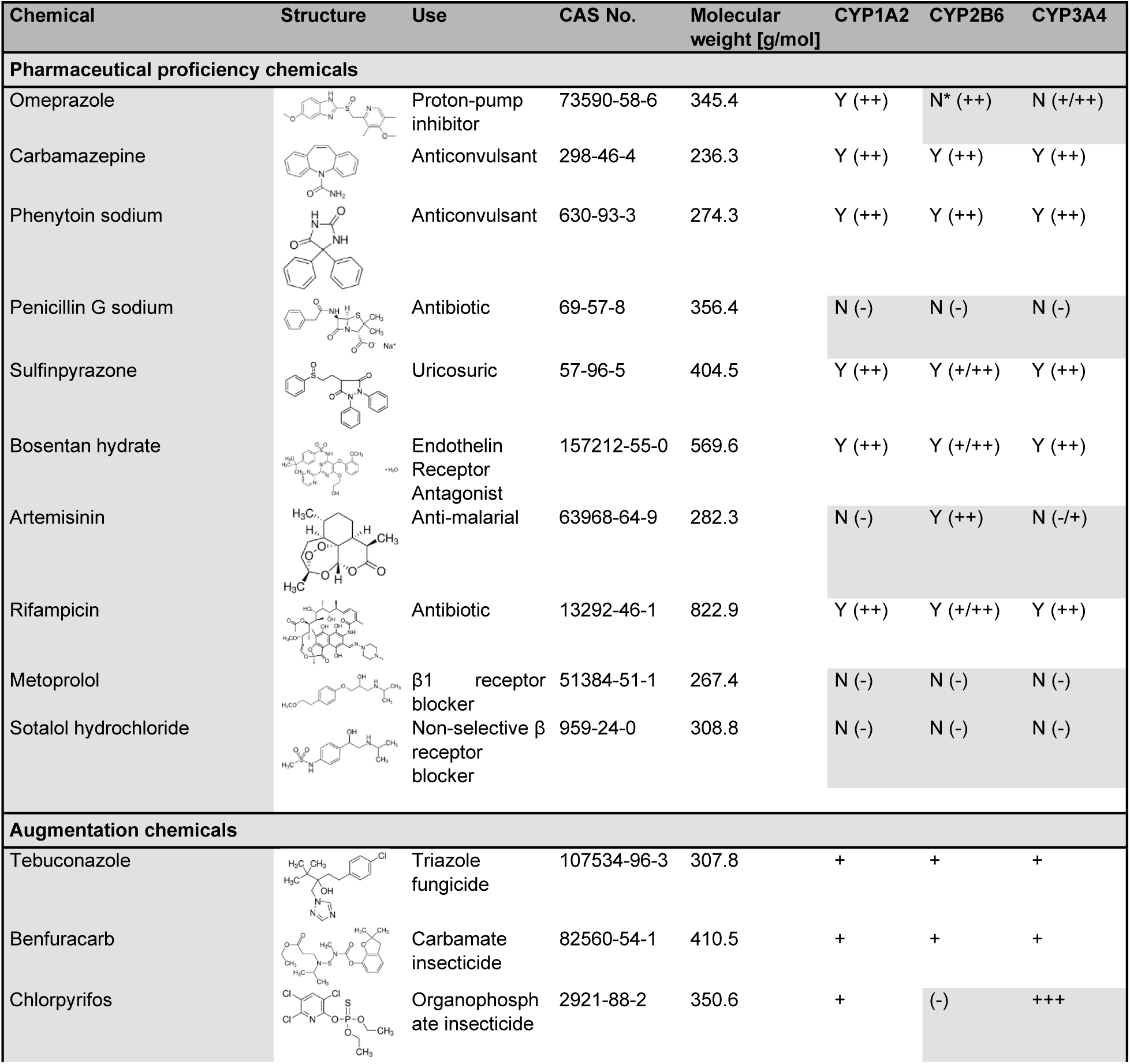
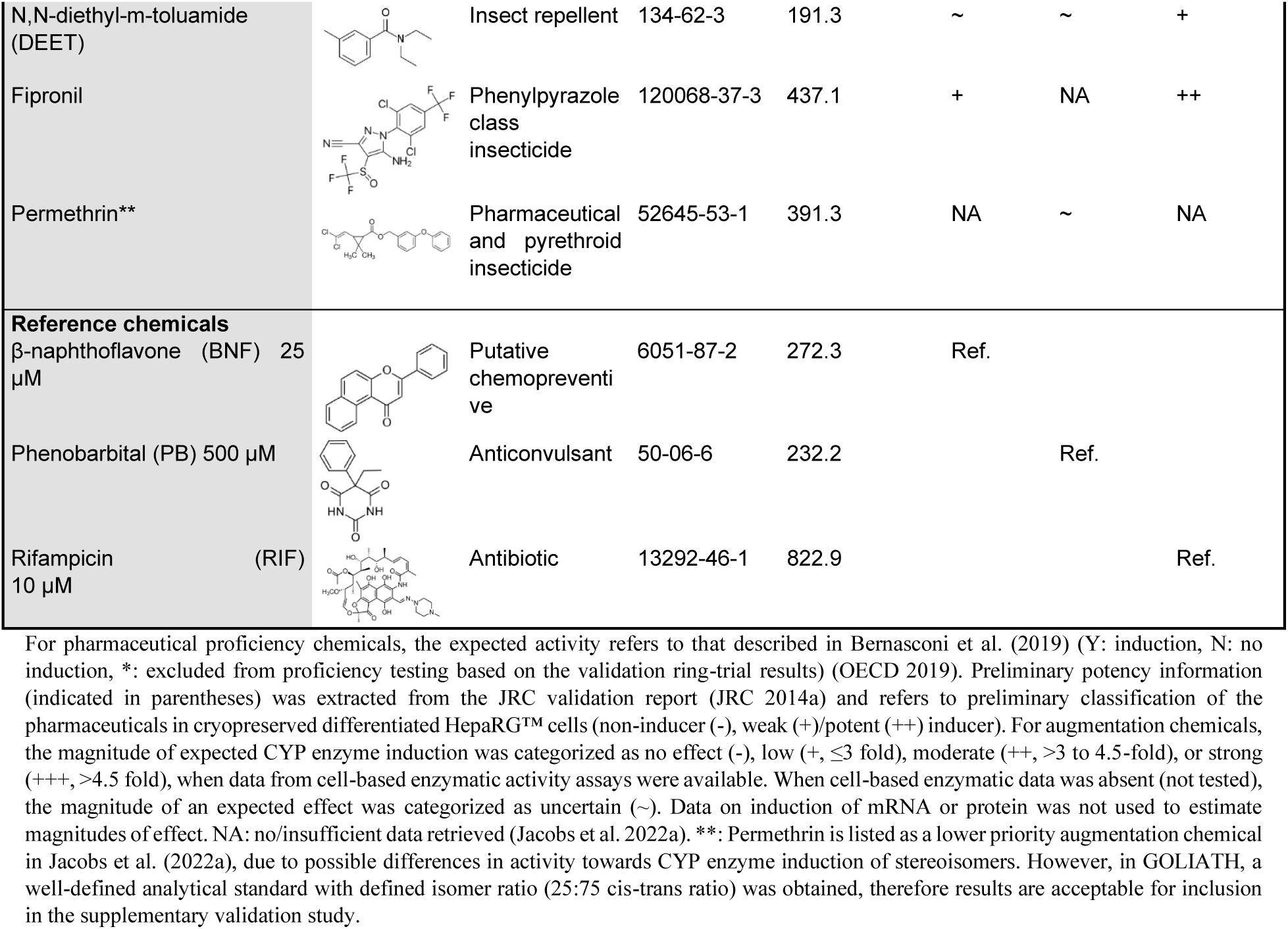
Chemicals tested during the GOLIATH supplementary chemical augmentation of the CYP enzyme induction test method and preliminary potency information. (as assessed in Jacobs et al. 2022a). Shaded cells exemplify induction/increase in catalytic activity.

The initial long list chemical set was first reviewed by two National Coordinators of the OECD Test Guidelines Programme, followed by independent peer review for scientific publication. Of these 23 candidate chemicals, 13 chemicals were proposed to augment the chemical applicability of the test method (Jacobs et al. 2022a). In line with the requirements agreed with the WNT, and also agreed within the GOLIATH grant agreement, 6 of the candidate chemicals were tested in two naïve laboratories to augment the chemical applicability domain: tebuconazole, benfuracarb, chlorpyrifos, N,N-diethyl-m-toluamide (DEET), fipronil and permethrin (Tab. 1, Augmentation chemicals). Both laboratories also tested the full set of 10 proficiency pharmaceutical chemicals (Tab. 1, Pharmaceutical proficiency chemicals) (Bernasconi et al. 2019), in order to demonstrate proficiency and successful method transfer and implementation prior to testing the 6 proposed augmentation chemicals. For the chemicals tested in the original JRC interlaboratory validation study (Bernasconi et al. 2019), preliminary potency information is included in the JRC validation study report (JRC 2014c).

The reference CYP positive inducers used and defined in the validation study (JRC 2014a, Bernasconi et al. 2019) were β-naphthoflavone (BNF) for CYP1A2, phenobarbital (PB) for CYP2B6, and rifampicin for the CYP3A subfamily (including CYP3A4).

An additional request from the WNT was for examples of regulatory applications of this test method. These have also been variously published (Jacobs et al. 2013; Bernasconi et al. 2019; Jacobs et al. 2022a), and are briefly discussed herein and the sister part 2 publication (Quartermain et al. 2026 submitted).

### 1.1 Rationale and general principle of the CYP Induction HepaRG™ Test Method

As previously described (e.g. Coecke et al. 2006, Jacobs et al. 2013), it is well established that acquiring relevant information on the metabolism of a substance is important when evaluating its toxic potential. The liver is usually the primary site of metabolism, but extra-hepatic tissues may also play a significant role. The enzymes that enable xenobiotic metabolism are traditionally divided into Phases I and II. The former involves oxidation of the parent molecule, e.g., by the mixed function oxidase activities of the many isoforms of the CYP family of enzymes. Phase II metabolism involves the conjugation of metabolites generated by Phase I oxidative reactions (or a conjugation with the parent chemical directly depending upon its structure), generally into more water soluble entities, with the most common Phase II reaction being glucuronidation and sulfation.

Most chemicals are metabolised via Phase I and Phase II reactions resulting in more polar forms that are partitioned to aqueous phases within the body and thus more readily excreted. Suitable *in vitro* assays can be used to gain information on the chemical fate of xenobiotics (parent substance and/or metabolite), which are ultimately responsible for an observed effect. In addition to xenobiotics, endogenous steroids are extensively metabolised by Phase I and II enzymes primarily in the liver but also in specific target tissues. Metabolism may lead to the inactivation of endogenous steroids and will affect the ability of the cells, and their tissues to perform their normal functions. Thus, exogenous chemicals that induce activity of Phase I and II enzymes can ultimately alter the availability of endogenous hormones and thereby potentially contribute to disruption of endocrine activity and possible endocrine disrupting effects.

Furthermore, one of the most frequently identified limitation of *in vitro* assays concerns the qualitative and quantitative deficiencies in the biotransformation of test chemicals in comparison with *in vivo* assays. These and related deficiencies are particularly important considerations when testing endocrine active substances (EAS), since several hormonally active chemicals, including some that occur naturally, are known to require bioactivation. The use of *in vitro* models to better prioritise chemicals for further testing, to advance the use of *in vitro* tests for directing testing that is more targeted for specific endpoints, and to move towards an eventual replacement of *in vivo* tests with *in vitro* are hampered by insufficient knowledge on chemical’s metabolism, bioactivation, and bioavailability.

The CYP induction test method aims to identify chemicals that trigger a CYP enzyme induction, which can be an MIE/early KE that may or may not have downstream toxic pathway consequences. It also aims to identify chemicals that, via CYP enzyme induction, may impact the biotransformation of endogenous as well as exogenous substances.

This test method can also be considered as an indicator of specific nuclear receptor activation, such as AhR (CYP1A1/1A2), PXR (CYP3A4) and CAR (CYP2B6). It is important to note however that for the test method’s current, validated status, it does not comprehensively assess full metabolism of the test chemical per se, nor general bioactivation or bioavailability but is specific only to endpoints of cytotoxicity and enzyme induction of CYP1A2, CYP2B6 and CYP3A4.

### 1.2 Study objective and test method purpose

The objective of this study was to establish the validated HepaRG™ CYP enzyme induction test method (JRC 2014a; Bernasconi et al. 2019) in two laboratories, and then demonstrate the applicability of the test method to non-pharmaceutical chemicals, including agrochemicals and industrial chemicals. For this purpose, a preliminary non-pharmaceutical chemical set (“augmentation chemicals”) was previously proposed (Jacobs et al. 2022a).

The purpose of this test method is to: “[have] reliable and relevant human hepatic *in vitro* metabolically competent test systems and transferable, reproducible and predictive *in vitro* methods to be used in integrated approaches for biotransformation and toxicological Mode of Action studies of substances and mixture/products of various industrial sectors.” (p. 5 of 164, JRC 2014a).

To achieve this objective two laboratories established the validated test method using the JRC Protocol (JRC 2014a, Bernasconi et al. 2019) with three reference chemicals and 10 pharmaceutical proficiency chemicals (Tab. 1). Upon successful completion, both laboratories tested the 6 proposed augmentation chemicals (Tab. 1). Next the within and between-laboratory reproducibility of the full proficiency chemical set (i.e., pharmaceutical and augmentation chemicals) was evaluated to derive the preliminary predictive capacity, in order to demonstrate the test method’s relevance in establishing CYP induction or not, and whether the chemical applicability domain for the proficiency chemicals could be expanded beyond the original pharmaceutical proficiency chemicals.

This paper describes the organisation, execution, results and conclusions of the chemical augmentation of the validation study conducted under the auspices of the EU funded GOLIATH project (Legler et al. 2019).

## 2 Materials and methods

### 2.1 Organisation of the chemical augmentation validation study and study design

The study was conducted according to the modular principles of validation and OECD Guidance Document 34, on the principles of conducting validation studies (Hartung et al. 2004, OECD 2005) as a ring trial with two laboratories. The trial design addressed the modules of test definition, within-laboratory reproducibility (WLR), transferability and between laboratory reproducibility (BLR). At this stage, and in agreement with the OECD WNT, as the test method had already been successfully validated previously (JRC 2014a, b, c; Bernasconi et al. 2019), two laboratories were considered sufficient to demonstrate the BLR of the 6 new proposed proficiency (‘augmentation’) chemicals.

One laboratory was technically proficient in metabolism studies (INRAE), and took the lead in setting up the test method in-house to confirm the SOP. The other laboratory (Utrecht University, UU) was naïve to the test method. All participants had extensive experience of maintaining mammalian cell culture and analytical chemistry including LC-MS/MS, but prior to this study, had no previous experience with this particular test method. Training was conducted online as on-site training could not be conducted due to limitations in international travel during the COVID-19 pandemic. Transfer to the naïve laboratory UU was conducted after INRAE had successfully established the method with the pharmaceutical proficiency chemicals (Person et al. 2026), INRAE were able to provide technical guidance to UU. Both laboratories demonstrated proficiency with the pharmaceutical chemicals, and then tested the augmentation chemical set of the same new proposed 6 selected proficiency chemicals.

Study management was coordinated by the UKHSA, the validation management study group consisted of UKHSA staff, the leads from INRAE and UU, and an independent validation expert and statistician. Non-blinded/uncoded test chemicals (6 non-pharmaceutical augmentation chemicals) were provided by a chemical repository coordinated at the Karolinska Institutet (KI). Prior chemical selection was conducted by UKHSA with independent expert review by two national coordinators to the OECD Test Guideline Programme and is reported elsewhere (Jacobs et al. 2022a).

Study management, provision of test chemicals via the chemical repository, and statistical analyses were conducted independently from the participating laboratories.

### 2.2 Chemicals and reagents

The following chemicals and reagents were used by both laboratories (Tab. 1), with the exception of metoprolol (CAS N°51384-51-1) (TRC, Canada MB38815 for UU and Sigma-Aldrich, COM448643953 for INRAE): Rifampicin (CAS N°13292-46-1), β-naphthoflavone (CAS N°6051-87-2), phenacetin (CAS N°62-44-2), bupropion HCl (CAS N°34911-55-2), acetaminophen (CAS N°103-90-2), formic acid (grade: LiChropur for LC-MS), DMSO (D2650, CAS N°67-68-5), (±)-hydroxybupropion-D6 (H062, CAS N°1184984-06-2), acetaminophen-D4 (P909, CAS N°64315-36-2), doxorubicin hydrochloride for cytotoxicity assessment (CAS N°25316-40-9), omeprazole (CAS N°73590-58-6), carbamazepin (CAS N°298-46-4), phenytoin sodium (CAS N°630-93-3), penicillin G sodium (CAS N°69-57-8), sulfinpyrazone (CAS N°57-96-5), bosentan hydrate (CAS N°157212-55-0), artemisinin (CAS N°63968-64-9), sotalol hydrochloride (CAS N°959-24-0), phenobarbital (CAS N°50-06-6) and midazolam HCl (CAS N°59467-70-8) (Sigma-Aldrich); hydroxybupropion (CAS N°357399-43-0), 1’-hydroxymidazolam (CAS N°59468-90-5) (Bertin Bioreagent); DPBS without Ca^2+^ and Mg^2+^ (PAN-Biotech); NaOH (VWR); Optima LC/MS-grade acetonitrile (≥ 99.9 %), LC-MS-grade methanol (≥ 99.9 %), micro-BCA Protein Assay Kit (#23235) (Thermo Fisher Scientific); and ultrapure water (Milli-Q system, Merck Millipore) was utilized for the preparation of HPLC mobile phases. Both participating laboratories purchased the reference chemicals, the enzymatic probe substrates and their respective metabolites, the internal standards and the pharmaceutical proficiency chemicals individually, but augmentation chemicals were supplied (non-blinded/ uncoded) via the independent central GOLIATH chemical repository, sourced from Sigma-Aldrich, (hosted at KI); both partners received an aliquot of the same chemical lot each. These were Tebuconazole (32013, purity 99.1 %), Benfuracarb (31554, purity 98.3 %) Chlorpyrifos (45395, purity 98.6 %) N,N-diethyl-m-toluamide (DEET) (36542, purity 98.5 %) Fipronil (46451, purity, ≥95 %) and Permethrin (cis:trans ratio 25:75, Y0001733, purity ≥95 %). Information related to all the chemicals tested is provided in Tab. 1.

For pharmaceutical proficiency chemicals, the expected activity refers to that described in Bernasconi et al. (2019) (Y: induction, N: no induction, *: excluded from proficiency testing based on the validation ring-trial results) (OECD 2019). Preliminary potency information (indicated in parentheses) was extracted from the JRC validation report (JRC 2014a) and refers to preliminary classification of the pharmaceuticals in cryopreserved differentiated HepaRG™ cells (non-inducer (-), weak (+)/potent (++) inducer). For augmentation chemicals, the magnitude of expected CYP enzyme induction was categorized as no effect (-), low (+, ≤3 fold), moderate (++, >3 to 4.5-fold), or strong (+++, >4.5 fold), when data from cell-based enzymatic activity assays were available. When cell-based enzymatic data was absent (not tested), the magnitude of an expected effect was categorized as uncertain (∼). Data on induction of mRNA or protein was not used to estimate magnitudes of effect. NA: no/insufficient data retrieved (Jacobs et al. 2022a). **: Permethrin is listed as a lower priority augmentation chemical in Jacobs et al. (2022a), due to possible differences in activity towards CYP enzyme induction of stereoisomers. However, in GOLIATH, a well-defined analytical standard with defined isomer ratio (25:75 cis-trans ratio) was obtained, therefore results are acceptable for inclusion in the supplementary validation study.

### 2.3 Cells and media

Cryopreserved HPR116 cells (HepaRG™ cells at their differentiated state at passage 16, HPR116-TA08 batches #253, #284, #295, #305, #312, #329 for biological replicates) were provided by Biopredic International (Saint-Grégoire, France). Manufacturer’s instructions were followed. Media used were HepaRG thaw, plate and general purpose medium with antibiotics (basal medium MIL600C [100 mL, 4°C] + Supplement ADD670C (14 mL, ࢤ20°C), HepaRG serum-free induction medium with antibiotics (basal medium MIL600C + supplement ADD650C [1.6 mL, ࢤ20°C]) from Biopredic International and incubation medium, Williams E without phenol red (500 mL, 4°C, Thermo Fisher Scientific A12176-01), HEPES 1M pH 7.4 (100 mL, 4°C, Thermo Fisher Scientific 15630-056), L-glutamine 200 mM (100 mL, ࢤ20°C, Thermo Fisher Scientific 25030-024).

### 2.4 Cell Culture Protocol

The SOP originally adhered to for this study was based on EURL ECVAM test method TM2009-14 (EURL ECVAM, 2009 <u>Cytochrome P450 (CYP) enzyme induction in vitro method using cryopreserved differentiated human HepaRG™ cells EURL</u> <u>ECVAM-TSAR</u>, last accessed 25 May 2026) and Bernasconi et al. (2019), with slight modifications and extension of its chemical applicability domain to include the augmentation chemicals specified in Tab. 1. It has also been reported in Person et al. (2026). Whilst the original SOP (EURL ECVAM 2009) was followed herein, it is not however the final adopted SOP for the OECD TG, as further small adaptations were made, as reported in part 2 of this study, Quartermain et al. 2026 submitted and published online in: https://tsar.jrc.ec.europa.eu/index.php/test-method/tm2009-14.

Briefly, HPR116 cells were thawed and seeded in monolayer at 7.2 × 10^4^ viable cells per well in a type I collagen coated 96-well plate (PLA136C, Biopredic International) and maintained in a humidified incubator at 37 °C with 5 % CO_2_. As recommended by the manufacturer, a cell viability after thawing above 80 % was required and confirmed by application of the trypan blue exclusion test. Six hours after seeding, cell morphology was observed and HepaRG thaw, plate and general-purpose medium was renewed for a further sixty-five hours, before exposure to test chemicals and running the test method. The time spent in the laminar airflow hood was minimized, and 96-well plates were kept on a preheated surface during this period, to reduce temperature fluctuations.

#### Cell Culture QA criteria and Validity of the Test

The seeded cryopreserved differentiated human HepaRG™ cells needed to meet the following quality control parameters:

- Cell viability after thawing: ≥80 % [e.g., by Trypan Blue exclusion test]
- Cell recovery per vial: ≥ 90 %
- Six hours after thawing, hepatocyte-like cells should be attached and appear in small, individual differentiated colonies
- After 72±0.9 h of culture, a restructuration of about 80 % confluent cryopreserved differentiated human HepaRG™ monolayer has to be observed with hepatocyte-like cells’ organization in clusters.

#### Validity of the experiment

For an experimental run to be valid, the following criteria were met:

1. Conduct tests in triplicate (i.e. 3 individual wells) for each test chemical concentration, reference chemical and solvent/vehicle control
2. The final solvent concentration during the CYP enzyme activity induction method should not affect cell viability and the induction of the specific isoform investigated (e.g., DMSO should not exceed 0.1 % v/v as DMSO induces CYP3A4)
3. The top concentration (e.g., 1 mM) should subsequently be soluble upon further dilutions in induction medium and should not be cytotoxic
4. Solvent/vehicle controls should demonstrate basal CYP enzyme activities in line with the data given on the test system/HPR116 cell batch) available from the suppliers Certificate of Analysis
5. Solvent/vehicle controls should demonstrate that the final concentration required has no effect on cell viability (must be within 15 % of the untreated controls)
6. Exposure to reference chemicals should lead to a ≥2-fold increase in CYP enzyme activity compared to solvent/vehicle control
7. At least six non cytotoxic test chemical concentrations have to be tested per chemical
8. At least two technical replicates available per test chemical concentration, reference chemical and solvent/vehicle control (in rare cases only two replicates were available, as e.g. there was loss during processing, or some aberrant data were identified by the laboratory and excluded)
9. At least three different thawed and independent cell batches must be used in independent runs.

### 2.5 Solubility and cytotoxicity assays

Solubility and cytotoxicity assays were performed for the 6 augmentation chemicals, by INRAE. The chemical repository, held at the Karolinska Institutet, Sweden (KI) also tested solubility as the chemicals arrived at the chemical repository. Solubility and cytotoxicity concentration ranges for the pharmaceutical proficiency chemicals were already available from the preceding validation study (Bernasconi et al. 2019 and JRC 2014a).

In a first step, the solubility in DMSO and in the treatment medium was assayed, as required in the test method’s specifications. At KI, solubility in DMSO was tested starting at 200 mM. All 6 augmentation chemicals were soluble at that concentration. INRAE started at 40 mg/mL in DMSO, and at 80 µg/mL in serum-free induction medium with DMSO (0.1 % v/v). The highest soluble concentration was further used as the starting concentration for cytotoxicity assessment. According to the cytotoxicity SOP provided by the JRC, cytotoxicity was assessed for 8 concentrations using a fluorescent method measuring the ability of living HPR116 cells to convert resazurin into resorufin (CellTiter-Blue® Cell Viability Assay, Promega, WI, USA).

### 2.6 Cytochrome P450 HepaRG induction test method

According to the TSAR SOP no.194 (2009), two test chemicals were assessed per plate, at six concentrations with a dilution factor of 1:2, starting from the highest soluble and non-cytotoxic concentration (Tab. 2).

**Tab. 2.** Concentrations of proficiency and augmentation chemicals used in this supplementary validation study. Concentrations for pharmaceutical proficiency chemicals were initially provided in the validation study (Bernasconi et al. 2019, JRC 2014a). For augmentation chemicals, testing concentrations were determined according to the SOP at INRAE and were followed by UU.

|  |  |  | Test conc.<br>[µg/mL] | Serial dilution test concentrations [µM] |  |  |  |  |  |
| --- | --- | --- | --- | --- | --- | --- | --- | --- | --- |
| Chemical | MW<br>[g/mol] | Solvent | 1 (highest) | 1 (highest) | 2 | 3 | 4 | 5 | 6 (lowest) |
| Original proficiency chemicals |  |  |  |  |  |  |  |  |  |
| Omeprazole | 345.4 | DMSO | 40.00 | 115.8 | 57.9 | 29.0 | 14.5 | 7.2 | 3.6 |
| Carbamazepine | 236.3 | DMSO | 40.00 | 169.3 | 84.6 | 42.3 | 21.2 | 10.6 | 5.3 |
| Phenytoin sodium | 274.3 | DMSO/<br>H<sub>2</sub>O (1:1) | 30.00 | 109.4 | 54.7 | 27.3 | 13.7 | 6.8 | 3.4 |
| Penicillin G sodium | 356.4 | DMSO | 40.00 | 112.2 | 56.1 | 28.1 | 14.0 | 7.0 | 3.5 |
| Sulfinpyrazone | 404.5 | DMSO | 40.00 | 98.9 | 49.4 | 24.7 | 12.4 | 6.2 | 3.1 |
| Bosentan hydrate | 569.6 | DMSO | 40.00 | 70.2 | 35.1 | 17.6 | 8.8 | 4.4 | 2.2 |
| Artemisinin | 282.3 | DMSO | 40.00 | 141.7 | 70.8 | 35.4 | 17.7 | 8.9 | 4.4 |
| Rifampicin | 822.9 | DMSO | 40.00 | 48.6 | 24.3 | 12.2 | 6.1 | 3.0 | 1.5 |
| Metoprolol | 267.4 | DMSO | 40.00 | 149.6 | 74.8 | 37.4 | 18.7 | 9.3 | 4.7 |
| Sotalol hydrochloride | 308.8 | DMSO | 40.00 | 129.5 | 64.8 | 32.4 | 16.2 | 8.1 | 4.0 |
| Augmentation Chemicals |  |  |  |  |  |  |  |  |  |
| Tebuconazole | 307.8 | DMSO | 40.00 | 130.0 | 65.0 | 32.5 | 16.2 | 8.1 | 4.1 |
| Benfuracarb | 410.5 | DMSO | 40.00 | 97.4 | 48.7 | 24.4 | 12.2 | 6.1 | 3.0 |
| Chlorpyrifos | 350.6 | DMSO | 40.00 | 114.1 | 57.0 | 28.5 | 14.3 | 7.1 | 3.6 |
| N,N-diethyl-m-toluamide (DEET) | 191.3 | DMSO | 40.00 | 209.1 | 104.5 | 52.3 | 26.1 | 13.1 | 6.5 |
| Fipronil | 437.1 | DMSO | 10.00 | 22.9 | 11.4 | 5.7 | 2.9 | 1.4 | 0.7 |
| Permethrin | 391.3 | DMSO | 10.00 | 25.6 | 12.8 | 6.4 | 3.2 | 1.6 | 0.8 |

Soluble, non-cytotoxic concentrations of augmentation chemicals were determined by INRAE. UU confirmed absence of test chemicals’ precipitation in DMSO and in treatment medium (visual observation)

The marker reactions of probe substrates were phenacetin O-deethylation (catalyzed by CYP1A2), bupropion hydroxylation (catalyzed by CYP2B6) and midazolam 1′-hydroxylation (catalyzed by CYP3A4). Briefly, 72 h after thawing, the assay was initiated by the addition of chemicals in serum-free induction medium (100 µL/well). Reference positive inducers β-naphthoflavone (CYP1A2, 25 µM), phenobarbital (CYP2B6, 500 µM) and rifampicin (CYP3A, 10 µM), as well as solvent controls (100 µL/well) were assessed on every plate. The final solvent concentration was 0.1 % (v/v) in “HepaRG serum-free induction medium”. The plate design included three technical replicates per condition. Outer wells were filled with DPBS.

Cells were cultured in serum-free conditions during the 48-hour exposure to test chemicals, reducing potential interferences and providing greater consistency. After 24 h of exposure, the exposure solutions were renewed and replaced by freshly prepared solutions for 24 h. At 48 h, exposure medium was removed, and all wells were washed twice with 100 µL incubation medium (Williams E without phenol red, HEPES 1M, L-glutamine 200 mM). A cocktail of the three CYP specific probe substrates (phenacetin, bupropion and midazolam) was prepared in a tube containing the methanol, wrapped with aluminum foil as bupropion is light-sensitive. It was evaporated under a stream of nitrogen at room temperature. The dried residue was reconstituted in 6 mL of incubation medium and vortexed to obtain the following final concentrations: phenacetin 26 µM, bupropion 100 µM and midazolam 3 µM. 50 µL of the substrate cocktail in incubation medium was added to each plate column with the same time-interval between columns. Substrate incubation was carried out during 60 ± 3 min at 37 °C in a humidified incubator with 5 % CO_2_. At the end of the incubation time, the reaction was quenched by stop solution containing (±)-hydroxybupropion-D6 (15 pg/µL) and acetaminophen-D4 (90 pg/µL) in ≥ 99.9 % acetonitrile, or (±)-hydroxybupropion-D6 (15 pg/µL) (UU only) in acetonitrile. 40 µL of cell medium were transferred to and mixed with 40 µL of ice-cold stop solution in a stop solution 96-well plate placed on ice.

### 2.7 Sample preparation

Calibration standards and quality controls were independently prepared by fortifying the incubation medium with intermediate solutions of probe metabolites in acetonitrile, as described in Person et al. (2026). For INRAE and UU: 40 µL of calibration standards, quality controls in matrix and blank and internal standard (ISTD) samples (incubation medium) were added into the outer wells of each stop solution plate and mixed with 40 µL stop solution at the same time as unknown samples, except for blank samples mixed with acetonitrile. This plate was centrifuged 10 min at 2200 x g, 4 °C. 30 µL of the particle-free supernatant was transferred into the corresponding wells of an analysis plate, thermo-sealed and stored at −20 °C for further LC-MS/MS analysis, which took place within a week. After thawing for analysis, 70 µL H_2_O was added and mixed directly in the plate (final acetonitrile content 15 % v/v in a total volume of 100 µL). The analysis plate was thermo-sealed and agitated for 20 s before introduction in a thermostated autosampler for further LC-MS/MS analysis.

### 2.8 LC-MS/MS analysis

Both laboratories used deuterated metabolites as internal standards, rather than DDIBA or griseofulvin (Bernasconi et al. 2019). The analytical (LC-MS/MS) method is not further specified in the Bernasconi et al. (2019) study, therefore this was qualified internally by each laboratory to meet the acceptance criteria for the probe substrates and their respective metabolites. Methodology further developed by INRAE and UU is described in detail by Person et al. (2026). The JRC validation study (Bernasconi et al. 2019) specified the lower limit of quantification (LLOQ) for the probe substrate metabolites as follows:

> “*LLOQ of 2.30 nM for acetaminophen, 1.15 nM for hydroxybupropion and 1.15 nM for 1-hydroxymidazolam were required before proceeding with sample analysis. Griseofulvin or 5.5-diethyl-1.3-diphenyl-2-iminobarbituric acid were included as reference items allowing correction for any loss of analyte during sample preparation and sample injection.*”

Although not entirely clear from this publication’s specification, it was assumed that these values were the final solutions required for sample injection. In this study, the limits of detection (LODs) and LLOQ achieved by the two laboratories for the different metabolites in cell culture medium (before water dilution) are presented in Tab. 3. n.d.: not determined. For instances where the lowest point of the calibration curve was reliably quantifiable, the LOD was not determined and the lowest point of the calibration curve is the LLOQ.

**Tab. 3.** Limits of detection and lower limits of quantification for the probe substate metabolites.

| Probe substrate metabolites |  | INRAE |  | UU |  |
| --- | --- | --- | --- | --- | --- |
|  |  | [nM] | [pg/µL] | [nM] | [pg/µL] |
| Acetaminophen | LOD | 10.3 | 1.56 | 3.0 | 0.46 |
|  | LLOQ | 31.0 | 4.67 | 9.0 | 1.36 |
| Hydroxybupropion | LOD | n.d. |  | 1.7 | 0.44 |
|  | LLOQ | 3.5 | 0.89 | 5.2 | 1.34 |
| 1'-Hydroxymidazolam | LOD | n.d. |  | n.d. |  |
|  | LLOQ | 3.3 | 1.11 | 1.3 | 0.44 |

n.d.: not determined. For instances where the lowest point of the calibration curve was reliably quantifiable, the LOD was not determined and the lowest point of the calibration curve is the LLOQ.

At INRAE, probe metabolites (acetaminophen, hydroxybupropion and 1′-hydroxymidazolam) were quantified by U-HPLC-MS/MS using a Nexera LC40 system coupled to a triple-quadrupole mass spectrometer (8045; Shimadzu, Kyoto, Japan). Chromatographic separation was achieved on a Phenomenex Synergi Polar-RP column (2.5 µm, 100 Å, 100 × 2.0 mm) maintained at 40 °C, with a binary mobile phase consisting of 0.1 % formic acid in water (A) and 0.1 % formic acid in methanol (B) at a flow rate of 0.35 mL/min. The gradient program was: 30 % B (0-0.5 min), 30-95 % B (0.5-2 min), 95 % B (2-3.5 min), 95-30 % B (3.5-3.6 min), and 30 % B (3.6-6 min).

Samples were injected (10 µL) from thermo-sealed 96-well plates maintained at 10 °C. Ionisation was performed using positive electrospray ionisation (ESI⁺), and metabolites were detected by multiple-reaction monitoring (MRM) with two transitions per analyte (for more details, see Person et al. 2026). Quantification was based on metabolite-area ratios relative to deuterated internal standards (acetaminophen-D₄ for acetaminophen and (±)-hydroxybupropion-D₆ for hydroxybupropion and 1’-hydroxymidazolam). Quality control samples at low, medium and high concentrations were included throughout each analytical run to ensure data integrity. SI-2 provides the LC-MS/MS SOP method, and the QC followed by INRAE, as an example.

At UU, a similar approach was taken, with slight modifications. Metabolites were quantified using a Shimadzu Nexera binary high-pressure LC coupled with Shimadzu MS/MS 8050 (Shimadzu, Kyoto, Japan). LC column was similar, with the exception that a Guard column C18 (4 x 2.0 mm ID) (Phenomenex) was used in combination with the RP column at 40 °C. The autosampler was maintained at 15°C and the injection volume was 1 μL. Optimal separation was achieved with a binary mobile phase (solvent A: 0.1 % formic acid in ultrapure water (milli-Q) and solvent B: 0.1 % formic acid in methanol) at a flow rate of 0.2 mL/min. The gradient elution program is: 0-0.5 min, 0 % B; 0.5-5.0 min, 0-95 % B; 5.0-7.0 min, 95 % B; 7.0-7.1 min, 95-0 % B; 7.1-10 min, 0 % B. The first 4.0 minutes the column eluent was diverted to waste. Needle wash solution consisted of ultrapure water/methanol (50:50, v/v). (More detailed information on the UU and INRAE concentrations of the probe substrates in the matrix are provided in SI-3, Tab. 1-3).

### 2.9 Protein content quantification

Residual incubation medium was aspirated off the cell plate and cells were lysed by the addition of 50 µL of 1 M sodium hydroxide and by mixing each column. The plate was covered and gently agitated for two hours at room temperature. Aliquots of the lysates were diluted with water (1:20) in duplicate, vortexed and stored at −20 °C until further analysis for protein content using the micro-BCA protein assay kit. Experiments were conducted as per the manufacturer’s instructions, consisting of a 2h-incubation at 37 °C followed by plate reading at a wavelength of 562 nm.

### 2.10 Data analysis

Each laboratory was responsible for checking the validity of the runs by applying the relevant validity of each criterion as specified in the SOP. Raw data of runs were entered by each study participant into a specifically prepared reporting template available via the TSAR (JRC TSAR 2009)/DB-ALM. The completed templates were checked and sent to the independent statistician for further checking, evaluation and statistical analysis as required.

After the validity check (see 2.4) of the submitted experiment data was confirmed, CYP activities were evaluated after protein content normalization and expressed in pmol of metabolite/min/mg of total protein. For each concentration and each CYP enzyme activity, fold-change was determined with respect to solvent controls. As suggested by Bernasconi et al. (2019), a ≥ 2-fold CYP enzyme activity was considered as an indication of CYP induction. Using this criterion, the experiment was considered:

- “Positive” for CYP induction, if ≥2-fold induction is observed for two or more consecutive test concentrations
- “Equivocal” for CYP induction, if ≥2-fold induction is observed at only one concentration, which is not the highest non-cytotoxic test concentration (with a plausible concentration-response curve)
- “Borderline” for CYP induction, if ≥2-fold induction is observed at only the highest test concentration (with a plausible concentration-response curve)
- “Negative” for CYP induction in all other cases.

Initially in alignment with Bernasconi et al. (2019), the results obtained were compared with known human CYP induction data (Bernasconi et al. 2019), and that obtained from the literature using human relevant cell systems (Jacobs et al. 2022a) by calculating the ratio between the maximum plasma concentration observed *in vivo* (Cmax) and the *in vitro* concentration required to produce a 2-fold induction response (F2) (Weiss and Haefeli, 2006; Kanebratt and Andersson, 2008; Grime et al., 2010). A Cmax/F2 ratio greater than 0.5 was used as the cutoff for predicting *in vivo* CYP induction. This threshold is considered conservative because it treats an *in vitro* concentration causing 2-fold induction as biologically relevant even when it is only half of the observed Cmax value. Although Cmax/EC50 ratios could also have been applied, EC50 values could not be determined for several of the validation and augmentation chemicals because complete concentration-response curves were unavailable.

As each chemical had to be tested in three different cell batches, an approach for obtaining a prediction across three experiments was needed. In the EURL ECVAM study, for *in vitro* classification, a chemical was considered a positive inducer overall if at least one experiment was positive, such that induction was detected in at least one batch of HepaRG cells across all three participating laboratories (using a two classifier/dichotomous approach). Whilst this was in line with US FDA guidance at the time (Bernasconi et al. 2019), the GOLIATH Validation Management Team re-examined this classification approach, considering the collective view of the ECVAM Scientific Advisory Committee (ESAC) and the results being generated within this study. This is further discussed in section 4.

Ultimately, the data of three valid experiments per chemical was interpreted following an updated multi-classifier data interpretation procedure (DIP). This was because the GOLIATH Validation Management Team were not confident that the revised classification criteria (i.e., dichotomous classification of “negative” or “positive” if two consecutive concentrations surpass 2-fold induction, or one out of three experiments/cell batches returns a “positive” call) sufficiently reflects the uncertainty associated with such borderline/equivocal test instances. We therefore proposed the following multi-classifier DIP to summarize the results across three experiments as described in Tab. 4.

**Tab. 4.** Decision tree for the conclusion of a test chemical (in three experiments/ three cell batches).

| Scenario | Exp. 1 | Exp. 2 | Exp. 3 | Conclusion |
| --- | --- | --- | --- | --- |
| 1 | positive | positive | positive | positive |
| 2 | 2 positive & 1 borderline, equivocal or negative |  |  | positive |
| 3 | 1 positive & 2 borderline |  |  | positive |
| 4 | 1 positive & 2 equivocal or negative (any combination) |  |  | equivocal |
| 5 | borderline | borderline | borderline | borderline |
| 6 | 2 borderline & 1 equivocal or negative |  |  | borderline |
| 7 | equivocal | equivocal | equivocal | equivocal |
| 8 | 2 equivocal & 1 borderline or negative |  |  | equivocal |
| 9 | 2 negative & 1 equivocal or borderline |  |  | negative |
| 10 | negative | negative | negative | negative |

Within-laboratory reproducibility (WLR) of a chemical was determined based on the concordance of valid experiment results. For each laboratory, the proportion of WLR chemicals among all chemicals was used as the main indicator of WLR. Potential reasons for a chemical not being WLR are discussed in section 3. The between-laboratory reproducibility (BLR) was determined as the proportion of chemicals with concordant results across the two laboratories (augmentation chemicals). The preliminary predictive capacity of the test method is described by the concordance of the results with the expected results as determined by the chemical selection.

### 2.11 Utilization of historical control data

Whilst it is important to achieve satisfactory performance in an experimental run, in line with historic data, it would be preferred if these historic data were established with the same [this] test method protocol to avoid confounding. It was noted that the protocol used to qualify cell batches by the supplier Biopredic International may differ from this protocol, e.g., in the duration of exposure to reference/probe substrates, concentrations, or the reference/probe substrates used. The study reported herein did not assess the extent to which these differences in the protocol may influence the measured enzymatic activity (i.e., basal or induced levels, in [pmol/min/mg protein]). Therefore, experiment-specific performance of the cell batches was compared with the batch-specific data provided by the supplier (see SI-4), and were considered satisfactory so were not used as an acceptance or exclusion criterion.

## 3 Results

The results are summarized below in tabular format. (Raw data including all analysis procedures and steps undertaken to transform the data are archived within the GOLIATH project and are available on request due to the extensive size of the files.)

### 3.1 Solubility and cytotoxicity results for the 6 augmentation chemicals

Results for the 6 augmentation chemicals assayed in three different cell batches for solubility and cytotoxicity conducted by INRAE, are provided in Tab. 5. The highest soluble concentration assayed in DMSO was 40 mg/mL. The highest soluble concentration in the medium, started from the 40 mg/mL stock solution in DMSO, to allow a cell treatment at the highest concentration of 40 µg/mL was 80 µg/mL (a 1:2 dilution factor was needed in the cytotoxicity assay) for Benfuracarb, Chlorpyrifos, DEET and Tebuconazole, or 20 µg/mL for Fipronil and Permethrin. Detailed cytotoxicity results and the corresponding graphs of three cell batches are provided in SI-1. Following cytotoxicity experiments, the highest concentration producing a cellular viability higher than 90 % after 48 h of incubation was used as the highest concentration for CYP induction assessment. Supplementary Information 1 (SI-1) provides the graphical representation for the solubility and cytotoxicity data for 6 augmentation chemicals together with SI Tab.1 cell viability data after thawing (INRAE).

**Tab. 5.** Summary of solubility and cytotoxicity results for the 6 augmentation chemicals.

| Augmentation chemicals | Highest concentration prepared in DMSO |  | Concentration prepared in medium |  | Highest non-cytotoxic concentration |  | Final solvent concentration (DMSO) |
| --- | --- | --- | --- | --- | --- | --- | --- |
|  | mg/mL | mM | µg/mL | µM | µg/mL | µM |  |
| Benfuracarb | 40 | 97.4 | 80 | 195 | 40.0 | 97.4 | 0.1 % |
| Chlorpyrifos | 40 | 114 | 80 | 228 | 40.0 | 114 | 0.1 % |
| Fipronil | 40 | 91.5 | 20 | 45.8 | 10.0 | 22.9 | 0.1 % |
| DEET | 40 | 209 | 80 | 418 | 40.0 | 209 | 0.1 % |
| Permethrin | 40 | 102 | 20 | 51.1 | 10.0 | 25.6 | 0.1 % |
| Tebuconazole | 40 | 130 | 80 | 260 | 40.0 | 130 | 0.1 % |

**Tab. 6.** Enzyme induction results of individual experiments with the pharmaceutical proficiency chemicals at INRAE and comparison with expected results from the JRC validation, with green indicating concordance and orange non-concordance. (+: positive; ࢤ: negative; E: equivocal; B: borderline; ‘: concentration-response not considered plausible; P: inducer; N: non-inducer; INC: inconclusive; *: inconclusive acc. to Tab. 4 and ‘P’ according to JRC (2014a); #: excluded from proficiency testing based on the validation ring trial results (Bernasconi et al. 2019; JRC 2014a); yellow highlights experiments with discordant results due to batch differences).

| Proficiency pharmaceuticals | Results (no. indicates cell batch) |  |  |  |  |  |  |  |  | Conclusion (as defined above) |  |  |
| --- | --- | --- | --- | --- | --- | --- | --- | --- | --- | --- | --- | --- |
|  | CYP1A2 |  |  | CYP2B6 |  |  | CYP3A4 |  |  | and comparison with expected result |  |  |
|  | 284 | 295 | 305 | 284 | 295 | 305 | 284 | 295 | 305 | CYP1A2 | CYP2B6 | CYP3A4 |
| Omeprazole | + | + | + | + | + | + | - | -' | - | P | P# | N |
| Carbamazepine | + | + | + | + | + | + | - | + | + | P | P | P |
| Phenytoin | + | + | + | + | + | + | + | + | + | P | P | P |
| Penicillin G | - | - | - | - | - | - | - | - | - | N | N | N |
| Sulfinpyrazone | + | + | + | + | + | B | + | + | + | P | P | P |
| Bosentan | + | + | + | - | - | - | + | + | + | P | N | P |
| Artemisinin | - | - | - | + | - | + | - | - | - | N | P | N |
| Rifampicin | + | + | + | E | - | + | + | + | + | P | INC/P* | P |
| Metoprolol | - | - | - | - | - | - | - | - | - | N | N | N |
| Sotalol | - | - | - | - | - | - | - | - | - | N | N | N |

Note: Omeprazole was classified as an *in vitro* inducer of CYP3A4 by all three laboratories testing HepaRG cells during the JRC validation study (JRC 2014a, Tab. M4.1, p. 101) but has not been reported to induce CYP3A *in vivo* clinically.

### 3.2 Test method transfer, proficiency, reproducibility and predictive capacity

#### 3.2.1 Method transfer

For the final data reported here, HPR116 cell batches were supplied to INRAE and UU by Biopredic International. Both laboratories purchased the reference chemicals, the enzymatic probe substrates and their respective metabolites, the internal standards and the pharmaceutical proficiency chemicals individually, but augmentation chemicals were supplied (non-blinded/ uncoded) via the independent central GOLIATH chemical repository (hosted at KI) that supported all the validation activities in the GOLIATH project; both partners each received an aliquot taken from the same chemical lot.

INRAE implemented the test method following the JRC SOP without further technical assistance; UU implemented the method under technical guidance from INRAE.

After qualification of the analytical chemistry method as set out in the JRC SOP and in line with guidance on the validation of bioanalytical methods (EMA 2022; FDA 2018; OECD 2018), both laboratories first demonstrated proficiency in successfully running the test method using the 10 previously validated pharmaceutical proficiency chemicals. Subsequently, they proceeded to test a subset of 6 of the 13 preliminary proposed non-pharmaceutical augmentation chemicals (Tab. 1).

#### 3.2.2 Proficiency of laboratories

Both laboratories tested the 10 pharmaceutical proficiency chemicals in three independent experiments using different cell batches, i.e., according to the validated JRC SOP (Bernasconi et al. 2019; JRC 2014a). Differences between the data interpretation procedure (DIP) of the JRC SOP and the revised DIP provided in section 2.10, were accounted for in the assessment of proficiency.

On the basis of an in-depth review (of each laboratory, experiment-by-experiment) of the data included in the JRC validation report (JRC 2014a), it was considered that achieving 100 % concordance with the pharmaceutical proficiency chemicals for all CYP isoforms is not realistic in practice. For example, data reported in Table M4.1 (p. 101 and 108 in the JRC validation report) indicate that omeprazole was identified as a CYP3A4 inducer in all three laboratories, and artemisinin was positive in one laboratory (CYP3A4) but was negative in two other laboratories). We therefore accepted proficiency when at least 80 % of the chemicals were classified concordantly, including the proficiency chemicals that are associated with less uncertainty and are known to be clear inducers of the CYP isoform in question. According to accepted OECD validation principles (OECD 2005), demonstration of successfully running and identifying all proficiency chemicals is required. But this does not mean that 100 % concordance is necessary.

#### 3.2.3 Evaluation of CYP induction reproducibility

Graphical representation of all HepaRG™ CYP induction data, the pharmaceutical proficiency chemicals and the 6 augmentation chemicals are provided in SI-5 and SI-6 respectively.

##### WLR proficiency pharmaceuticals results for each participating laboratory

The WLR results for INRAE are provided in Tab.6, according to the new multi-classifier DIP (note that this was first reported in Person et al. (2026), according to the old JRC (2014a) DIP). As indicated by the green shading (three columns on the far right), all conclusions for CYP1A2 and CYP3A4 were concordant with the expected results as provided by JRC (2014a) and Bernasconi et al. (2019). In all these cases the new DIP defined here and the two classifier previous DIP utilized in JRC (2014a) and Bernasconi et al. (2019) resulted in the same conclusion. The CYP2B6 conclusions for the 9 proficiency chemicals with expected results was concordant in 8 instances (88.9 %). The only exception was bosentan, which did not induce CYP2B6, although it was expected to. For rifampicin, the conclusions of the two data interpretation procedures differed. In summary, it was concluded that INRAE successfully demonstrated proficiency for all three CYP isoforms.

Analyzing the data regarding WLR, i.e., concordance of results across experiments per CYP, revealed that overall, in 26 of 30 cases, i.e., 86.7 %, the three experiments had concordant results, however reproducibility differed between CYPs, with 100 % WLR for CYP1A2, 90 % for CYP3A4 and 70 % WLR for CYP2B6. However, the differences between CYP2B6 experiments observed for sulfinpyrazone, artemisinin and rifampicin were not very clear, as curve-shapes were similar, or a threshold effect (for rifampicin) was observed (Person et al. 2026). In contrast, the first test chemical experiment with carbamazepine in relation to CYP3A4 induction was clearly different from the other two experiments, where a clear concentration-response and induction up to 10-fold was reached (see SI-5).

As indicated by the green shading (three columns on the far right), all conclusions for CYP1A2 and CYP3A4 were concordant with the expected results as provided by JRC (2014a) and Bernasconi et al. (2019). In all these cases the new DIP defined here and the two classifier DIP utilised in JRC (2014a) and Bernasconi et al. (2019) resulted in the same conclusion. The CYP2B6 conclusions for the 9 proficiency chemicals with expected results was concordant in 8 instances (88.9 %). The only exception was bosentan, which did not induce CYP2B6, although it was expected to. For rifampicin, the conclusions of the two data interpretation procedures differed. In summary, it was concluded that INRAE successfully demonstrated proficiency for all three CYP isoforms.

Tab. 7 summarizes the results for UU.

**Tab. 7:** Enzyme induction results of individual experiments with the proficiency chemicals at UU and comparison with expected results from the JRC validation, with green indicating concordance and orange non-concordance. (+: positive; ࢤ: negative; E: equivocal; B: borderline; ‘: concentration-response not considered plausible; P: inducer; N: non-inducer; INC: inconclusive; *: inconclusive acc. to Tab. 4 and ‘P’ according to the draft TG; **: borderline acc. to Tab. 4 and ‘N’ according to the draft TG #: excluded from proficiency testing based on the validation ring trial results (JRC 2014a); yellow highlights experiments with discordant results due to batch differences).

| Proficiency chemicals | Results (no. indicates cell batch) |  |  |  |  |  |  |  |  | Conclusion (defined above) |  |  |
| --- | --- | --- | --- | --- | --- | --- | --- | --- | --- | --- | --- | --- |
|  | CYP1A2 |  |  | CYP2B6 |  |  | CYP3A4 |  |  | and comparison with expected result |  |  |
|  | 312 | 253 | 329 | 312 | 253 | 329 | 312 | 253 | 329 | CYP1A2 | CYP2B6 | CYP3A4 |
| Omeprazole | + | + | + | + | + | + | + | - | - | P | P# | INC/P* |
| Carbamazepine | + | + | + | + | + | + | + | + | + | P | P | P |
| Phenytoin | + | + | + | + | + | + | + | + | + | P | P | P |
| Penicillin G | - | - | - | + | + | - | - | - | - | N | P | N |
| Sulfinpyrazone | + | + | + | + | + | + | + | + | + | P | P | P |
| Bosentan | + | + | + | + | E | E | + | + | + | P | INC/P* | P |
| Artemisinin | - | - | - | E | - | + | - | - | - | N | INC/P* | N |
| Rifampicin | + | + | + | + | + | E | + | + | + | P | P | P |
| Metoprolol | B | - | B | + | - | + | + | - | - | B/N** | P | INC/P* |
| Sotalol | - | - | - | - | - | + | - | - | - | N | INC/P* | N |

Note as for Tab. 6: Omeprazole was classified as an *in vitro* inducer of CYP3A4 by all three laboratories testing HepaRG cells during the JRC validation study (JRC 2014a, Tab. M4.1, p. 101) but has not been reported to induce CYP3A *in vivo* clinically.

As indicated by the green shading, all conclusions for CYP1A2 were concordant with the expected results as defined in the JRC validation (2014a). The CYP2B6 conclusions were correct for 6 of the 9 proficiency chemicals with expected results. Penicillin G, metoprolol and sotalol were incorrectly concluded to be inducers of CYP2B6 (i.e., false-positives). CYP3A4 results were in agreement with the expected results for 8 proficiency chemicals. Omeprazole and metoprolol were incorrectly predicted as inducers. In six instances the two DIPs led to different conclusions. In five instances the two classifier procedure would have resulted in positive rather than ‘inconclusive’ classifications, which reflects the less conservative, binary data classification interpretation approach initially suggested (JRC 2014a). In summary, it was concluded that UU demonstrated satisfactory proficiency for CYP isoforms 1A2 and 3A4 (≥ 80 % concordance), but lower accuracy was noted regarding the identification of CYP2B6 inducing chemicals.

Analyzing the data regarding WLR, concordance of results across experiments per CYP, revealed that overall, this occurred for 21 of 30 cases (70.0 %). Again, the limited data available suggested that reproducibility may differ between CYPs, with 90 % WLR for CYP1A2, 40 % for CYP2B6 and 80 % WLR for CYP3A4. The concentration-response curves for CYP1A2 with metoprolol were almost identical between experiments - the difference being introduced by threshold only. Also, the single positive CYP3A4 experiments for omeprazole and metoprolol were not very strongly positive, with maximum inductions of 2.1-fold and 2.34-fold, respectively (see SI-5). For CYP2B6, a clear difference between experiments was observed for Penicillin G and metoprolol. Interestingly, concentration-response curves of bosentan, artemisinin and rifampicin consistently showed a decrease in induction with increasing concentrations - a trend also reflected by results reported by INRAE (see SI-5) and similar to observations in the JRC validation study (JRC 2014a). Differences between experiments were observed at the lowest test concentrations. Finally, the discordant results for sotalol were a threshold effect. Whilst a bit less evident than the results generated by INRAE, a more detailed evaluation of the concentration-response curves does support the conclusion of CYP-specific WLR.

##### WLR of the 6 augmentation chemicals results for each participating laboratory

After demonstrating proficiency, both laboratories tested the 6 augmentation chemicals. Concentration response curve data are available graphically in supplementary information, and a summary is presented in Tab. 8.

**Tab. 8.** INRAE and UU enzyme induction results of individual experiments with the augmentation chemicals and comparison with expected literature derived results, with green indicating concordance and orange non-concordance. (+: positive; ࢤ: negative; E: equivocal; B: borderline; P: inducer; N: non-inducer; INC: inconclusive; *: inconclusive according to Tab. 4 and ‘P’ according to the JRC (2014a); **: borderline acc. to Tab. 4 and ‘N’ according to JRC (2014a); greyed text: expected results unknown).

| INRAE Results for augmentation chemicals |  |  |  |  |  |  |  |  |  |  |  |  |
| --- | --- | --- | --- | --- | --- | --- | --- | --- | --- | --- | --- | --- |
|  | Results (no. indicates cell batch) |  |  |  |  |  |  |  |  | Conclusion (defined above) |  |  |
|  | CYP1A2 |  |  | CYP2B6 |  |  | CYP3A4 |  |  | and comparison with expected result |  |  |
|  | 253 | 312 | 329 | 253 | 312 | 329 | 253 | 312 | 329 | CYP1A2 | CYP2B6 | CYP3A4 |
| Permethrin | - | - | - | B | B | B | - | - | - | N | B/N** | N |
| Fipronil | + | + | + | + | + | E | + | + | + | P | P | P |
| Tebuconazole | + | + | E | + | + | + | + | + | + | P | P | P |
| DEET | + | + | + | + | + | - | + | + | + | P | P | P |
| Chlorpyrifos | B | - | - | - | - | - | B | - | B | N | N | B/N** |
| Benfuracarb | + | - | B | + | + | + | + | B | B | INC/P* | P | P |
| UU Results for augmentation chemicals |  |  |  |  |  |  |  |  |  |  |  |  |
| Permethrin | - | - | - | - | + | B | - | - | - | N | INC/P | N |
| Fipronil | + | + | + | - | + | E | + | + | + | P | INC/P | P |
| Tebuconazole | E | + | + | E | + | + | E | + | + | P | P | P |
| DEET | E | + | + | - | + | E | + | + | + | P | INC/P* | P |
| Chlorpyrifos | B | B | B | - | - | - | - | + | - | B/N** | N | INC/P |
| Benfuracarb | - | + | + | + | + | + | B | + | + | P | P | P |

The WLR at UU was 8/18 = 44.4 %. Excluding the results obtained with the cell batch no. 253 results in concordance of the two remaining experiments of 14/18 = 77.8 %.

At INRAE the WLR was 11/18 = 61.1 %. In most instances, the concentration-response curves were similar, and differences were due to the threshold. The most pronounced difference between experiments was observed for DEET for CYP2B6.

##### BLR of the 6 augmentation chemicals results for each participating laboratory

For INRAE, all conclusions for CYP1A2 and CYP3A4 were concordant with the expected results as defined from the original validation study for pharmaceutical proficiency chemicals. In all these cases the DIP defined here and the initial one, came to the same conclusion. The CYP2B6 conclusions for the 9 proficiency chemicals with expected results were correct in 8 instances (88.9 %). The single exception was bosentan, which did not induce CYP2B6, although it was expected to. For rifampicin, the conclusions of the two data interpretation procedures differed. In summary, it was concluded that INRAE successfully demonstrated proficiency for all three CYP isoforms.

For UU, the CYP2B6 conclusions were correct for 6 of the 9 proficiency chemicals with expected results. Penicillin G, metoprolol and sotalol were incorrectly concluded to be inducers of CYP2B6 (i.e., false-positive). CYP3A4 results agreed with the expected results for 8 proficiency chemicals. Omeprazole and metoprolol were incorrectly predicted as inducers. As noted in section 2.2, metoprolol was sourced differently from INRAE, and this may contribute to the difference seen. In 6 cases the two data interpretation procedures led to different results. In five of these instances, the procedure defined here would have resulted in ‘inconclusive’, which reflects the conservative DIP approach described in the JRC validation report (2014a) and Bernasconi et al. (2019). In summary, it was concluded that UU demonstrated satisfactory proficiency for CYP isoforms 1A2 and 3A4 (≥ 80 % concordance), but higher uncertainty was noted regarding the identification of CYP2B6 inducing chemicals. Higher uncertainty for CYP2B6 was also noted in the JRC validation.

#### 3.2.4 Predictive capacity

The analysis of the predictivity of the test method for the augmentation chemicals was limited by two aspects. First, for two chemicals (permethrin and fipronil) expected results could not be defined for some CYP isoforms. Second, the 6 chemicals were expected to induce all CYPs, except chlorpyrifos, which was expected to be a non-inducer of CYP2B6. However, both laboratories correctly identified the CYP-inducing properties with 12/15 = 80 % (INRAE) and 14/15 = 93 % (UU) accuracy. At INRAE, two borderline conclusions (permethrin: CYP2B6; chlorpyrifos: CYP3A4) were observed, which translate into ‘non-inducer’ according to the JRC (2014a) interpretation. Both cases were inconclusive at UU, but interpreted as ‘inducer’, due to one positive experiment. Both laboratories observed no or very weak CYP1A2 induction with chlorpyrifos, resulting in a negative conclusion, but the expected result was positive.

## 4. Discussion

Metabolism is multifactorial and occurs over distinct phases. Several human xenobiotic metabolising enzymes are inducible by pharmaceuticals and environmental chemicals. Phase I CYP enzyme induction can have an impact on the biological effect(s) of a given xenobiotic and/or on endogenous physiological functions and signalling pathways. Lasting modulation of the functionality of CYP enzymes may have a profound impact on general metabolism, since these key enzymes, in addition to the biotransformation of xenobiotics, are involved in numerous biochemical reactions with key roles in energy metabolism, including the anabolism and catabolism of sugars, lipids and hormones. Several CYPs are involved in anabolic processes, and a large number of steroids derived from cholesterol, may see their concentration impacted upon by an induction of CYPs. For instance, the first step of cholesterol biotransformation in this context is known to be impacted by some dioxins.

Indeed, CYP induction per se, following the nuclear receptor-xenobiotic interaction, has been suggested as an important biological event in several Adverse Outcome Pathways (AOPs) for over 15 years (Pelkonen et al. 2008; USEPA, 2011; Jacobs et al. 2013, Vinken et al. 2013, also discussed in JRC 2014a). It has also been explored in relation to the identification of metabolic disruption (Person et al. 2026). Indeed, the HepaRG™ CYP induction test method will be applicable for the investigation of the MIEs and very early KEs of most complex human health endpoints as part of an Integrated Approach to the Testing and Assessment (IATA) of chemicals. Also, by contributing to the mechanistic Phase I metabolism understanding, it will support the paradigm transition from *in vivo* rodent studies, to human relevant *in vitro* test batteries.

Having been previously successfully validated with pharmaceutical proficiency chemicals (Bernasconi et al. 2019; JRC 2014a; JRC 2014b; JRC 2014c), the HepaRG™ CYP induction test method reported can address a core aspect of CYP induction, specifically in relation to key CYPs of interest in drug and xenobiotic metabolism with probe substrates for CYP1A2 and the AhR, CYP3A4 and PXR, and CYP2B6 and CAR. It can elucidate the ability of a test chemical to induce these hepatic enzymes, but is not currently validated to identify test chemical metabolites. Inclusion of metabolism relevant *in vitro* test methods is an identified need for regulatory applications and Integrated Approaches for the Testing and Assesment of chemicals (OECD 2008; Bernasconi et al. 2019; Jacobs et al. 2013; 2022a). It specifically generates human relevant information on CAR/PXR activation, and can therefore contribute to support and refine chemical safety assessment for toxicological modes of action where species differences are often not quantified.

Whilst at this point in time,approaches for evaluating *in vitro* metabolism cannot be used to qualitatively or quantitatively predict (de)toxification that may occur *in vivo,* they can provide valuable insights and contribute to weight of evidence mechanistic analyses.

To date, this has been the only test method proposed to the OECD Test Guideline Programme and that has then been fully validated, that addresses human *in vitro* metabolism via induction of selected CYP enzymes. Other CYP relevant test methods known to be utlised for regulatory purposes include the ethoxyresorufin-O-demethylase (EROD) assay has been used as a measure of CYP1A(1) induction in standardized tests (e.g., with fish: ISO/TS 23893-2:2007(en)), and an AhR CALUX test method for environmental matrices has been undergoing validation at the International Standards Organisation (ISO)). Neither of these are as comprehensively relevant to human CYP mechanistic understanding, as the validated HepaRG™ CYP induction test method.

The analysis of the predictivity of the test method for the augmentation chemicals was limited by two aspects. First, for two chemicals (permethrin and fipronil), expected results could not be defined for some CYP isoforms. Second, the 6 chemicals were expected to induce all CYPs, except for chlorpyrifos, which was expected to not to be a CYP2B6 inducer. Both laboratories correctly identified the CYP-inducing properties with 12/15 = 80 % (INRAE) and 14/15 = 93 % (UU) accuracy. At INRAE, two borderline conclusions (permethrin: CYP2B6; chlorpyrifos: CYP3A4) were observed, which translate into ‘non-inducer’ according to the two classifier interpretation. Both cases were inconclusive at UU, but were interpreted as ‘inducer’, according to the two classifier DIP, due to one positive experiment. Both laboratories observed no or very weak CYP1A2 induction with chlorpyrifos, resulting in a negative conclusion, whilst the expected result was positive.

The interpretation of the results is further complicated by the fact that the experiment run with cell batch no. 253 consistently gave the lowest response in all UU experiments. While the effect on the predictivity was limited, the WLR was substantially affected. As INRAE and UU tested the same cell batches with the augmentation chemicals and did not see such batch-specific effects with cell batch 253, and this was also not seen in the historical data provided by Biopredic International (SI-4), this is most likely an artifact of relatively higher basal enzyme activities in the solvent control, resulting in relatively lower fold-induction. Nevertheless, the respective experiment with cell batch 253 passed acceptance criteria and was therefore not excluded from the analysis.

Between batch and between laboratory variability is inevitable to a certain degree, therefore assessment of ‘sporadic’ cases should be considered. The practical relevance of a consistent positive finding in one batch (or perhaps two) in one test facility amongst otherwise negative or irregular concentration-response relationships, needed to be considered. At the time of conducting this work, the recommendations in the FDA guidance (2020) were that if the chemical results in an induction according to preset criteria in one out of three batches of primary hepatocytes, the chemical is regarded as an inducer and a clinical DDI study is needed. However, this is a consideration developed for pharmaceutical safety and efficacy assessment, as opposed to chemical hazard assessment, where regulatory decisions require a substantive weight of evidence basis upon which to conclude. With respect to chemical hazard assessments and the use of TGs, the primary objective is to characterize the intrinsic toxicological properties of the test chemical itself for the endpoint of concern, rather than characterizing effects on toxicodynamic parameters that will influence the safety profile of pharmaceuticals.

Uncertainty from variable data could be addressed by defining an appropriate statistical measure to support the call on a test concentration/an experiment e.g. exceeding the 2-fold induction threshold and with statistical significance, as for example implemented for *in vitro* steroidogenesis (OECD TG 456, 2025).

A ‘sporadic’ finding was with an assumed negative proficiency chemical metoprolol. Understanding whether the one positive consistent curve (one batch, one facility) for one activity (CYP2B6) was a reliable result, needed further batch testing, which was not conducted at the time. If the chemical is a new test item under pharmaceutical development, the 2020 FDA recommendation is that: an *in vivo* investigation is required, and this is also consistent with the European Medicines Agency recommendation (EMA 2024). However, it is not a sufficient evidence base upon which to trigger *in vivo* investigation for industrial chemicals. Further limitations in regulatory applications of this test method as currently validated include the assessment of inhibition of CYPs, Phase II metabolism induction/inhibition and the identification of metabolites generated by test chemicals. These are all development aspects that this test method could in future be modified for, and validated.

With the generation of additional chemical augmentation data in two laboratories, within the GOLIATH project, it was hoped that this work would be adequate to support the further peer review and TG adoption at the OECD. However, in light of the discussion here, as the performance of the second laboratory (UU) was not as robust as that for INRAE and opportunities to consolidate the data by testing more HPR116 cell batches at UU were not possible at that time, at the conclusion of the experimental work reported here, this final objective was not quite ready to pursue. To meet this objective, further funding to support additional naïve laboratory work was sought by the OECD project lead, the successful outcome of which is reported in part 2 (Quartermain et al. 2026).

The robustness of the test method as a tool for the assessment of (semi)quantitative conclusions on chemicals’ CYP induction potential could be further strengthened by anchoring not only to the solvent-control, but also to the signal strength obtained for the positive reference inducer for each CYP isoform. Currently, the acceptance criterion for a valid experiment is that the reference inducer results in ≥2-fold induction of the target CYP isoform, the same level that is required for an (unknown) test chemical to be classified as an inducer. In cases with unusually greater basal enzyme activity, or lower induction with the reference chemical, this decreases the dynamic range of classifying a chemical as a (non-)inducer, without rendering the experiment invalid. However, one would expect the reference inducer concentration to be set at a level which allows some dynamic range for the classification of a chemical as an inducer - particularly as in the case of inducible CYP isoforms there is no clear upper bound level of induction. At the stage of the experimental work conducted herein, the reference and proficiency data generated are not sufficient yet to inform on a test chemical’s potency to induce a certain CYP isoform, and that a potency-based tripartite (or more granular) classification is therefore not supported yet. Nevertheless, it would be desirable if in the future, as this test method is used and generates data, it could be developed to provide potency data. In due time, this could be proposed as a retrospective data analysis exercise, to further strengthen the test method’s utility for chemical safety assessment. We are confident that such anchoring to the positive reference inducer, and thus a step towards more quantitative information on induction potency, will become possible once more experience with this test method is gained and control data are generated. This will help characterise the dynamic range between cell batches, and generate a historic control database not only for basal enzyme activity levels, but also for the positive reference chemicals. Moving beyond the initial binary approach taken for the original validation, concentration-response data has been provided (Tab. 7 and 8; SI-5 and 6), and as more data are generated this will be collectively useful for broadening regulatory applications towards the development of *in silico* tools and IATA applications (Jacobs et al. 2022b).

## 5. Conclusions and recommendations

The supplementary chemical augmentation validation of the HepaRG CYP enzyme induction test method was successfully conducted and completed, with promising results. Additional aspects for refinement, such as definition of criteria to inform upon a chemicals’ potency or activity bands are recommended. In addition, further recommendations to improve the robustness of the data interpretation procedure, especially with respect to the identification of borderline/weak CYP enzyme inducing chemicals have been identified applying a multi-classifier approach.

Overall, this test method was successfully established in two laboratories, both of which demonstrated proficiency by correctly identifying CYP (non-)inducing pharmaceutical chemicals for isoforms 1A2 and 3A4, and partially for CYP2B6 (one laboratory with some uncertainty). Subsequently, the test method’s applicability towards non-pharmaceutical chemicals, including pesticides and industrial chemicals was demonstrated, thereby closely meeting a key requirement of the OECD TGP requested in 2019 and one of the objectives of the OECD TGP workplan project no. 4.76.

These considerations are further explored in a second, follow up study conducted at UKHSA laboratories, and are reported in Part 2 (Quartermain et al). Taken together, parts 1 and 2, these data were sufficient for the OECD peer review to progress towards the drafting of the HepaRG CYP induction TG, which was adopted in late April 2026.

## Disclaimer

The views and opinions expressed in this article do not represent the official position of the authors respective institutions, or of the European Commission.

## Acknowledgements

Foremost, the authors wish to thank Sandra Coecke and Camilla Bernasconi at the EC Joint Research Centre, Ispra, Italy for their leading work in coordinating the validation of this test method on the basis of *in vivo* human pharmaceutical data. The authors also gratefully acknowledge Christophe Chesne, CEO of Biopredic International, for his guidance to this work.

## Supplementary material attached Data Availability

All additional data is available in Supplementary information.

## Declaration of generative AI and AI-assisted technologies in the manuscript preparation process

Microsoft copilot was used to formulate the materials and methods section in the ALTEX style.

