## Supplemental information for "Chemical augmentation of the validated HepaRG^TM^ CYP enzyme induction test method Part 1: The Goliath two laboratory study"

*Jacobs et al*

**Supplementary information 1.**

**Graphical presentation of the cytotoxicity results for the 6 augmentation chemicals**

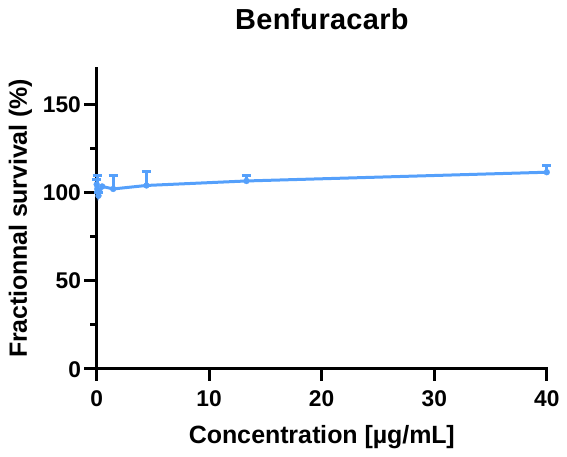

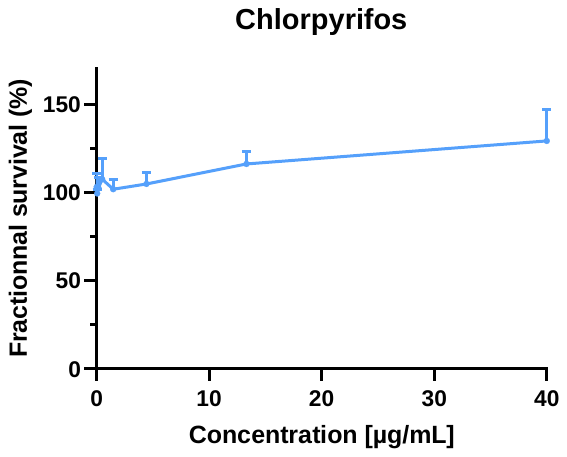

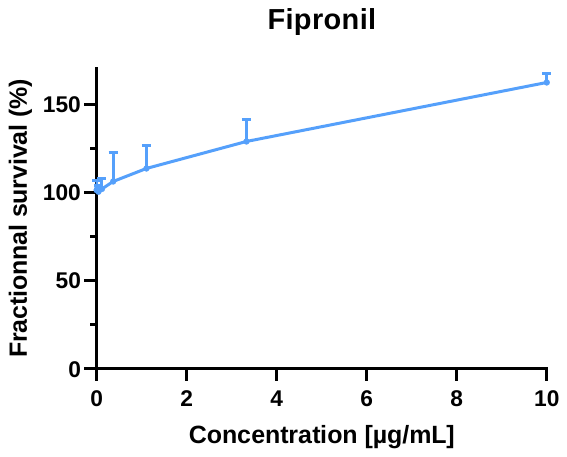

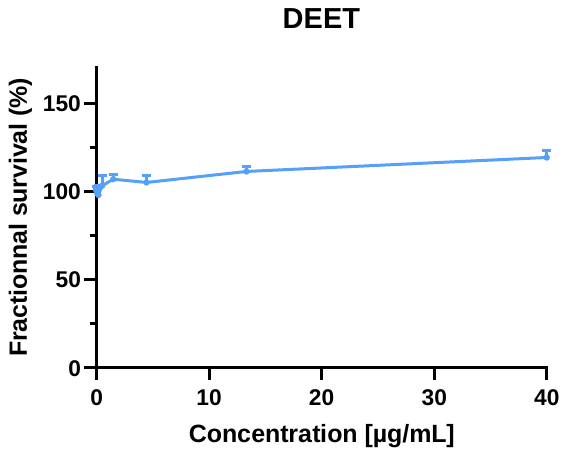

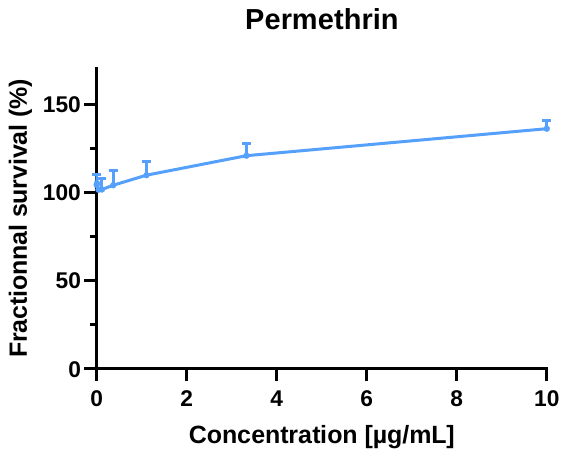

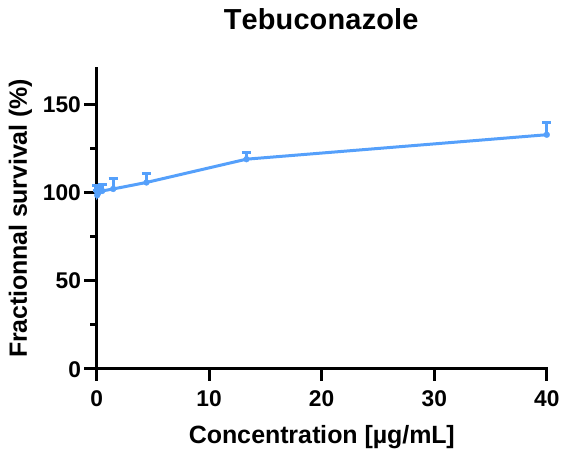

Detailed cytotoxicity results for the 6 augmentation chemicals (mean ± SD of three HPR116 cell batches)

Tab. 1. Cell viability after thawing. Data provided by INRAE

| **Cell batches** | **Cell viability after thawing (%)** |
| --- | --- |
| 253 | **91** |
| 284 | **92** |
|  | **91** |
|  | **86** |
|  | **86** |
| 295 | **84** |
|  | **86** |
|  | **89** |
| 305 | **88** |
|  | **90** |
|  | **88** |
|  | **86** |
| 312 | **90** |
|  | **81** |
| 329 | **81** |

**Supplementary Information 2**

The HepaRG CYP induction assay SOP used in this study is the same as that published (JRC2014a and Bernasconi et al 2019) so is not reproduced here. With the original SOP, guidance was not provided for the LC-MS/MS aspects, as instrumentation varies greatly.

The methods followed by INRAE for LC-MS/MS are provided here

The production of specific metabolites (probe substrates) by CYP1A2, CYP2B6 and CYP3A4 is quantified by LC-MS/MS analysis.

STOCK AND WORKING SOLUTIONS

Stock solutions of the three metabolites 1’-Hydroxymidazolam (OH-MID), Hydroxybupropion (OH-BUP), and Acetaminophen (ACETA) are prepared at 700 mg/L (metabolites) in glass vials:

- OH-MID and OH-BUP are weighed (minimum 1 mg) and solubilized in methanol
- ACETA is weighed (minimum 1 mg) and solubilized in acetonitrile

(*Excel file for concentration calculations*)

- Stock solutions are vortexed 5 min on a multiple tube shaker Multi Reax (speed 10)
- Metabolite mix and following intermediate dilutions in acetonitrile are performed in duplicate (for CS and QC independent preparation) in acetonitrile from stock solutions in microtubes, as described in Tables 3 and 4.

**Tab. 1. Metabolite mix preparation in acetonitrile (prepared in duplicate)**

|  | **Metabolite Mix** | | | |
| --- | --- | --- | --- | --- |
|  | **[C] (pg/µL)** | **Stock solution (µL)** | **ACN (µL)** | **Final volume (µL)** |
| **OH-MID** | 175 000 | 250 | 200 | 1 000 |
| **OH-BUP** | 140 000 | 200 |  |  |
| **ACETA** | 245 000 | 350 |  |  |

**Tab.2. Intermediate dilutions in acetonitrile (prepared in duplicate)**

| **Intermediate dilutions** | | | | | | |
| --- | --- | --- | --- | --- | --- | --- |
| **Dilution N°** | **Added volume (µL)** | **From solution** | **ACN volume (µL)** | **Final volume (µL)** | **Concentration (pg/µL)** | |
| **P1** | 100 | Metabolite Mix | 200 | 300 | **OH-MID** | 58 333.3 |
|  |  |  |  |  | **OH-BUP** | 46 666.7 |
|  |  |  |  |  | **ACETA** | 81 666.7 |
| **P2** | 150 | P1 | 50 | 200 | **OH-MID** | 43 750.0 |
|  |  |  |  |  | **OH-BUP** | 35 000.0 |
|  |  |  |  |  | **ACETA** | 61 250.0 |
| **P3** | 150 | P2 | 75 | 225 | **OH-MID** | 29 166.7 |
|  |  |  |  |  | **OH-BUP** | 23 333.3 |
|  |  |  |  |  | **ACETA** | 40 833.3 |
| **P4** | 150 | P3 | 200 | 350 | **OH-MID** | 12 500.0 |
|  |  |  |  |  | **OH-BUP** | 10 000.0 |
|  |  |  |  |  | **ACETA** | 17 500.0 |
| **P5** | 50 | P4 | 200 | 250 | **OH-MID** | 2 500.0 |
|  |  |  |  |  | **OH-BUP** | 2 000.0 |
|  |  |  |  |  | **ACETA** | 3 500.0 |
| **P6** | 50 | P5 | 200 | 250 | **OH-MID** | 500.0 |
|  |  |  |  |  | **OH-BUP** | 400.0 |
|  |  |  |  |  | **ACETA** | 700.0 |
| **P7** | 50 | P6 | 100 | 150 | **OH-MID** | 166.7 |
|  |  |  |  |  | **OH-BUP** | 133.3 |
|  |  |  |  |  | **ACETA** | 233.3 |
| **P8** | 50 | P7 | 100 | 150 | **OH-MID** | 55.6 |
|  |  |  |  |  | **OH-BUP** | 44.4 |
|  |  |  |  |  | **ACETA** | 77.8 |

CALIBRATION STANDARDS, QC, BLANKS AND STOP SOLUTIONS

Calibration standards (CS) are prepared by fortifying the matrix (Incubation Medium) with intermediate solutions of metabolites in acetonitrile, as presented in Table 5.

**Tab. 3. Calibration standards preparation in incubation medium**

| **Calibration standards (CS)** | | | | | | |
| --- | --- | --- | --- | --- | --- | --- |
| **N°** | **Added volume (µL)** | **From solution** | **Incubation medium (µL)** | **Final volume (µL)** | **Concentration (pg/µL)** | |
| **CS1** | 5 | P1 | 245 | 250 | **OH-MID** | 1 166.7 |
|  |  |  |  |  | **OH-BUP** | 933.3 |
|  |  |  |  |  | **ACETA** | 1 633.3 |
| **CS2** | 5 | P2 | 245 | 250 | **OH-MID** | 875.0 |
|  |  |  |  |  | **OH-BUP** | 700.0 |
|  |  |  |  |  | **ACETA** | 1 225.0 |
| **CS3** | 5 | P3 | 245 | 250 | **OH-MID** | 583.3 |
|  |  |  |  |  | **OH-BUP** | 466.7 |
|  |  |  |  |  | **ACETA** | 816.7 |
| **CS4** | 5 | P4 | 245 | 250 | **OH-MID** | 250.0 |
|  |  |  |  |  | **OH-BUP** | 200.0 |
|  |  |  |  |  | **ACETA** | 350.0 |
| **CS5** | 5 | P5 | 245 | 250 | **OH-MID** | 50.0 |
|  |  |  |  |  | **OH-BUP** | 40.0 |
|  |  |  |  |  | **ACETA** | 70.0 |
| **CS6** | 5 | P6 | 245 | 250 | **OH-MID** | 10.0 |
|  |  |  |  |  | **OH-BUP** | 8.0 |
|  |  |  |  |  | **ACETA** | 14.0 |
| **CS7** | 5 | P7 | 245 | 250 | **OH-MID** | 3.33 |
|  |  |  |  |  | **OH-BUP** | 2.67 |
|  |  |  |  |  | **ACETA** | 4.67 |
| **CS8** | 5 | P8 | 245 | 250 | **OH-MID** | 1.11 |
|  |  |  |  |  | **OH-BUP** | 0.89 |
|  |  |  |  |  | **ACETA** | 1.56 |

Quality Controls (QC) are independently prepared in the matrix from independent intermediate solutions of metabolites in acetonitrile, using the dilution scheme in Table 4.

**Tab. 4. QCs preparation in incubation medium in triplicate**

| **QC preparation** | | | | | | |
| --- | --- | --- | --- | --- | --- | --- |
| **N°** | **Added volume (µL)** | **From solution** | **Incubation medium (µL)** | **Final volume (µL)** | **Concentration (pg/µL)** | |
| **QC1-High** | 10 | P2 | 490 | 500 | **OH-MID** | 875.00 |
|  |  |  |  |  | **OH-BUP** | 700.00 |
|  |  |  |  |  | **ACETA** | 1 225.00 |
| **QC2-Mid** | 10 | P3 | 490 | 500 | **OH-MID** | 583.33 |
|  |  |  |  |  | **OH-BUP** | 466.67 |
|  |  |  |  |  | **ACETA** | 816.67 |
| **QC3-Low** | 10 | P7 | 490 | 500 | **OH-MID** | 3.33 |
|  |  |  |  |  | **OH-BUP** | 2.67 |
|  |  |  |  |  | **ACETA** | 4.67 |

CS and QC are prepared for each analysis plate and extracted at the same time as unknown samples.

40 µL of CS and QC in matrix is added directly into the “Stop solution calibration plate” (*cf*. Figure 5.)

Blank samples are prepared by adding 40 µL of incubation medium to 40 µL of acetonitrile directly into the “Stop solution calibration plate”

ISTD blank sample is prepared by adding 40 µL of incubation medium directly into the “Stop solution calibration plate”

Ice-cold stop solution with (±)-hydroxybupropion-D6 (15 pg/µL in acetonitrile) and acetaminophen-D4 (90 pg/µL in acetonitrile) is prepared and 40 µL is added in the Stop solution calibration plate CS, ISTD blank, QC and samples wells (quenching). Final concentrations are 7.5 pg/µL and 45 pg/µL respectively.

An example of an analytical plate is presented Figure 1.

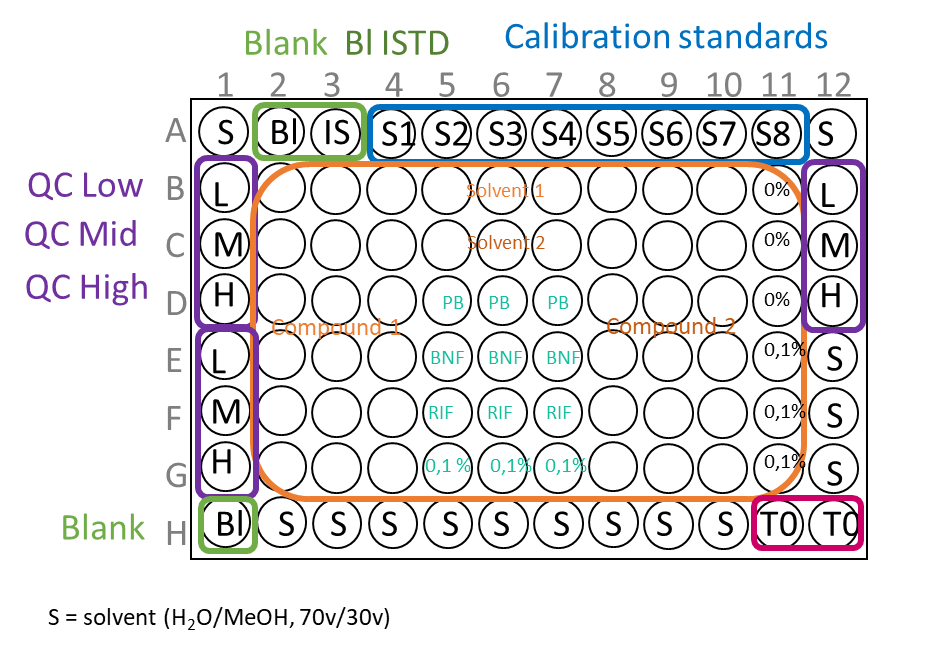

*Figure 1. Plate layout for LC-MS/MS analysis*

LC-MS/MS METHOD - CHROMATOGRAPHIC SETTINGS

Reversed-phase high-performance liquid chromatographic separation (RP-HPLC) is achieved on an Shimadzu Nexera system. Metabolites are separated using a Phenomenex Synergi 2.5 µm polar-rp 100a 100 mm ø 2.0 mm thermostated at 40°C. The autosampler is maintained at 10°C and the injection volume is 10 μL from 96-well plate covered with peelable heat sealing foil. Optimal separation is achieved with a binary mobile phase (solvent A: 0.1 % formic acid in water and solvent B: 0.1 % formic acid in methanol) at a flow rate of 0.35 mL/min. The gradient elution program is: 0-0.5 min, 30 % B; 0.5-2 min, 30-95 % B; 2-3.5 min, 95 % B; 3.5-3.6 min, 95-30 % B; 3.6-6 min, 30 % B.

MASS SPECTROMETRY SETTINGS

Metabolites are detected and quantified using a triple quadrupole mass spectrometer Shimadzu 8045 equipped with a heated ElectroSpray Ionisation source (ESI) in positive mode. An investigation of optimal collision energies was conducted for each fragmentation and two transitions were selected: quantification and confirmation (Tab. 5).

**Tab. 5. Mass spectrometry parameters for the detection of the metabolites**

| **Chemical substrates** | **Precursor ion *(m/z)*** | **Product ions *(m/z)*** | | **Dwell** | **Collision energy (V)** | | **Retention time (min)** |
| --- | --- | --- | --- | --- | --- | --- | --- |
|  |  | ***Quantification*** | ***Confirmation*** | **(ms)** | ***Quantification*** | ***Confirmation*** |  |
| Acetaminophen | 152.2 | 110 | 65.05 / 93.05 | 22 | -18 | -31/-23 | 1.47 |
| Hydroxybupropion | 256.2 | 139.05 | 238.1 / 131.1 | 22 | -26 | -13/-27 | 2.45 |
| 1’-Hydroxymidazolam | 342.2 | 324.1 | 203.1 / 168.15 | 22 | -20 | -27/-28 | 2.91 |

Optimal parameters are: nebulizing gas flow: 3 L/min, heating gas flow: 10 L/min, interface temperature: 300°C, DL temperature: 250°C, heat block temperature: 400°C, drying gas flow: 10 L/min.

*Figure 7. Plate layout for protein determination*

**Supplementary Information 3**

**Analytical Chemistry method: Concentrations of CYP probe substrates in the matrix for UU and INRAE**

**Tab.1 Acetaminophen:**

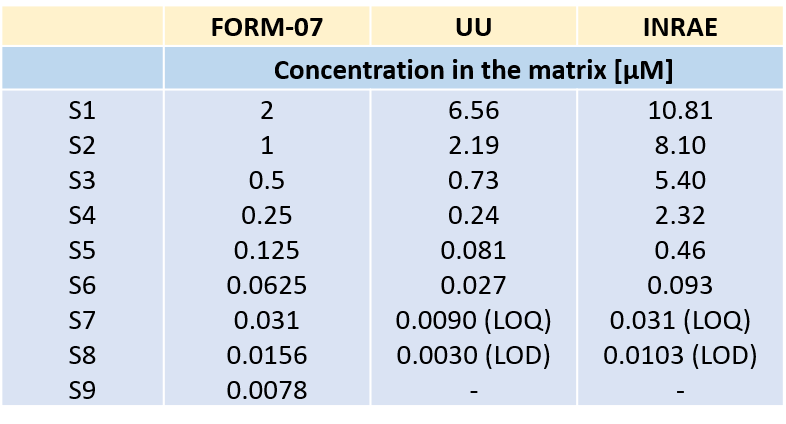

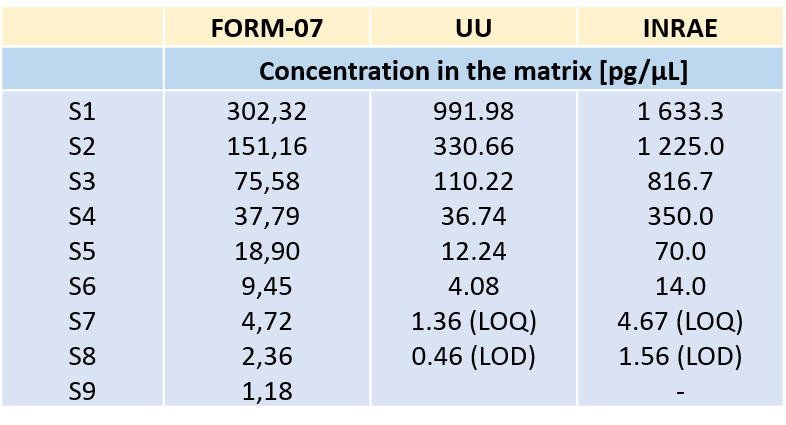

**Tab.2 Hydroxy-bupropion**

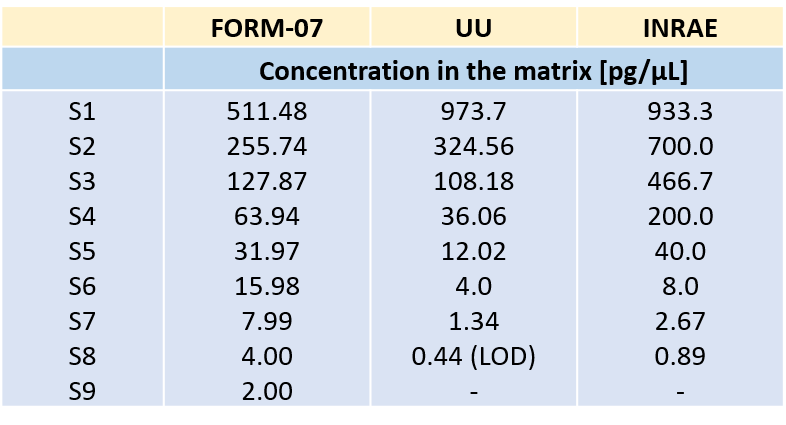

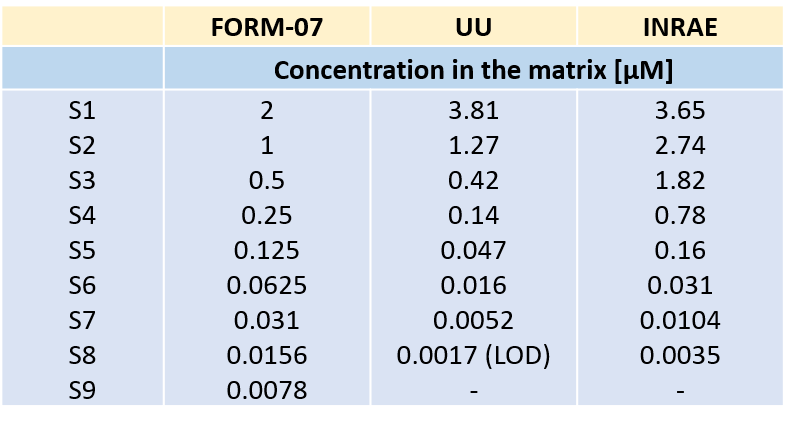

**Tab 3 Hydroxy-midazolam**

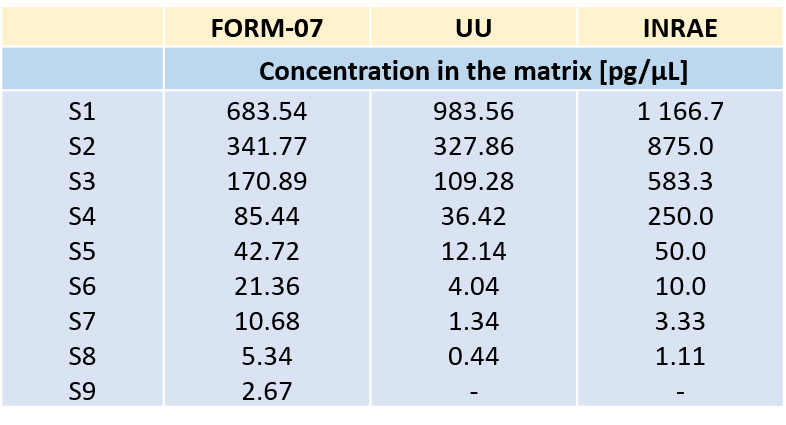

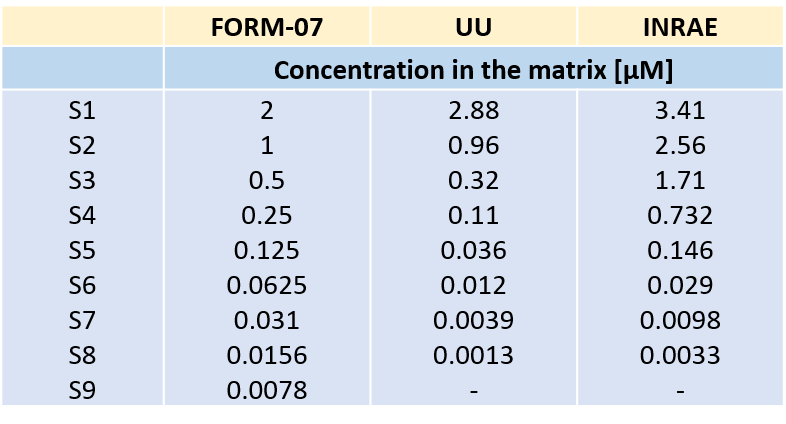

**Supplementary Information 4**

**Historic QC data for specific HPR116 cell batches provided by Biopredic International**

For QC purposes, BPI has records on historical performance of **170 batches of HPR116 cells**, recorded between 2010 – 2022.

Tab. 1 Summary of the HPR116 CYP induction QC protocol at BPI

| **Timeline** | **Experimental step description** |
| --- | --- |
| Day 1/ 0 h | thawing cells (Thawing/Plating/General Purpose medium 670, 10% FCS, 0.5% DMSO) |
| Day 1/ 6 h | observe and document (if possible) cell morphology: phase contrast light microscopy |
| Day 4/ 72 h | replace general purpose medium with HepaRG induction medium 640 (2% FCS, 0.5% DMSO) with test **chemicals** (*here: inducer probe chemicals*) |
| Day 5/ 96 h | renew HepaRG induction medium 640 with test chemicals |
| Day 6/ 120 h | end of incubation with test chemicals;  incubation of cells with test **substrates** |

Tab. 2 QC procedure and conditions at BPI

|  | **CYP1A2** | **CYP2B6** | **CYP3A4/5** |
| --- | --- | --- | --- |
| Inducer probe chemical | Omeprazole 50 µM | Phenobarbital 1 mM | Rifampicin 10 µM |
| Test substrate | Phenacetin | Bupropion | Midazolam |

Table 12: Historic data of HPR116 CYP induction QC at BPI (170 batches, 2010-2022)

|  | Basal levels  [pmol metabolite/ mg protein/ min] | | | Induced levels  [pmol metabolite/ mg protein/ min] | | | Fold induction (no unit) | | |
| --- | --- | --- | --- | --- | --- | --- | --- | --- | --- |
|  | **CYP1A2** | **CYP2B6** | **CYP3A4/5** | **CYP1A2** | **CYP2B6** | **CYP3A4/5** | **CYP1A2** | **CYP2B6** | **CYP3A4/5** |
| **Q25** | 0.20 | 0.10 | 0.21 | 1.10 | 0.40 | 2.30 | 4.0 | 3.6 | 8.3 |
| **Median** | 0.26 | 0.14 | 0.38 | 1.46 | 0.62 | 4.08 | 5.2 | 4.5 | 10.8 |
| **Q75** | 0.33 | 0.18 | 0.53 | 1.70 | 0.93 | 5.06 | 6.7 | 5.8 | 12.9 |

Tab.3 Historic QC data obtained at BPI for specific batches that were shared with GOLIATH partners basal and induced level are given in [pmol metabolite/ mg protein/ min], while fold-induction has no unit.

|  |  |  | **CYP1A2** | | | **CYP2B6** | | | **CYP3A4/5** | | |
| --- | --- | --- | --- | --- | --- | --- | --- | --- | --- | --- | --- |
| **Batch #** | **GOLIATH partner** | **prod. year** | **basal level** | **induced level** | **fold-ind.** | **basal level** | **induced level** | **fold-ind.** | **basal level** | **induced level** | **fold-ind.** |
| #284 | INRAE | 2018 | 0.22 | 1.64 | 7.6 | 0.15 | 0.8 | 5.4 | 0.40 | 4.3 | 10.8 |
| #295 | INRAE | 2019 | 0.22 | 1.58 | 7.1 | 0.21 | 0.7 | 3.2 | 0.73 | 8.4 | 11.5 |
| #305 | INRAE | 2019 | 0.21 | 0.80 | 3.8 | 0.14 | 0.3 | 2.3 | 0.27 | 2.3 | 8.6 |
| #253 | UU | 2018 | 0.25 | 1.86 | 7.4 | 0.11 | 0.4 | 4.0 | 0.49 | 4.0 | 8.2 |
| #312 | UU | 2020 | 0.51 | 1.79 | 3.5 | 0.26 | 0.5 | 2.0 | 0.53 | 2.7 | 5.1 |
| #329 | UU | 2021 | 0.29 | 1.79 | 6.2 | 0.29 | 1.2 | 4.0 | 0.76 | 7.2 | 9.5 |

**Supplementary information 5**

**Graphical representation of the results for the pharmaceutical proficiency chemicals**

Note: y-axes ranges are not aligned

**Omeprazole**

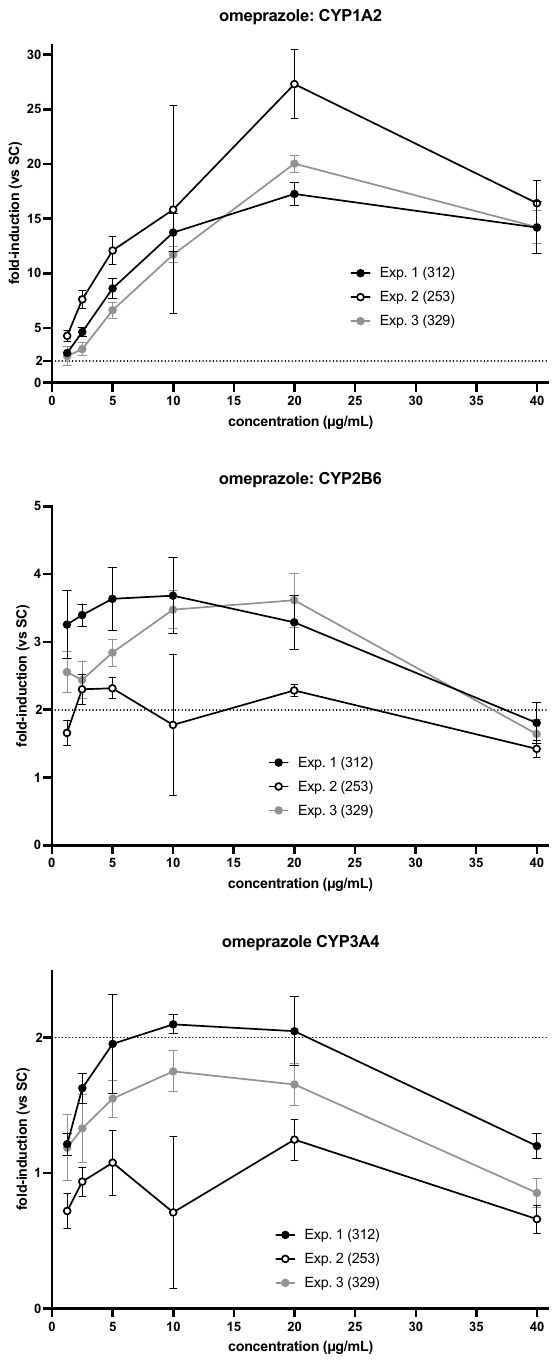
**INRAE	UU**

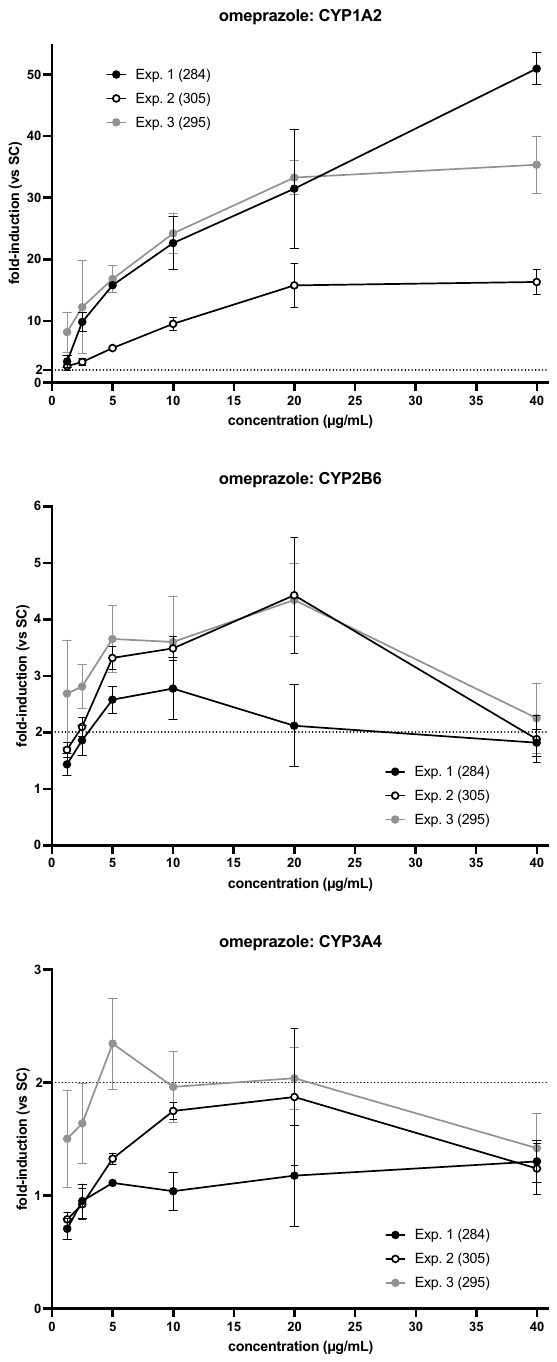

**Carbamazepine**

**INRAE	UU**

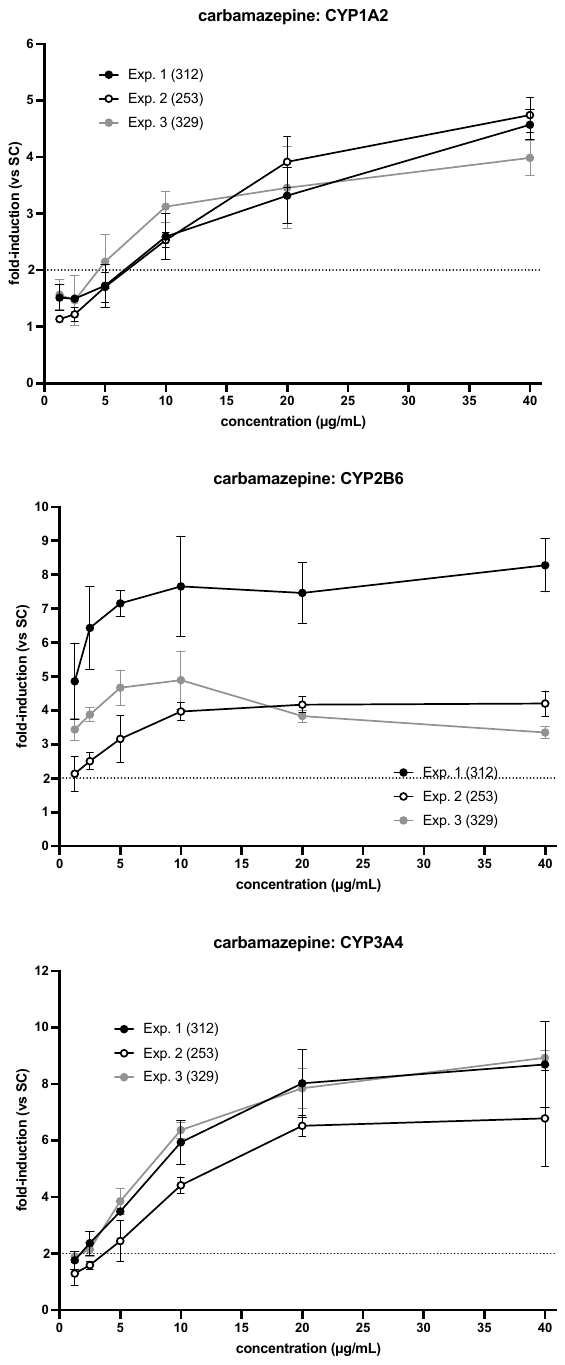

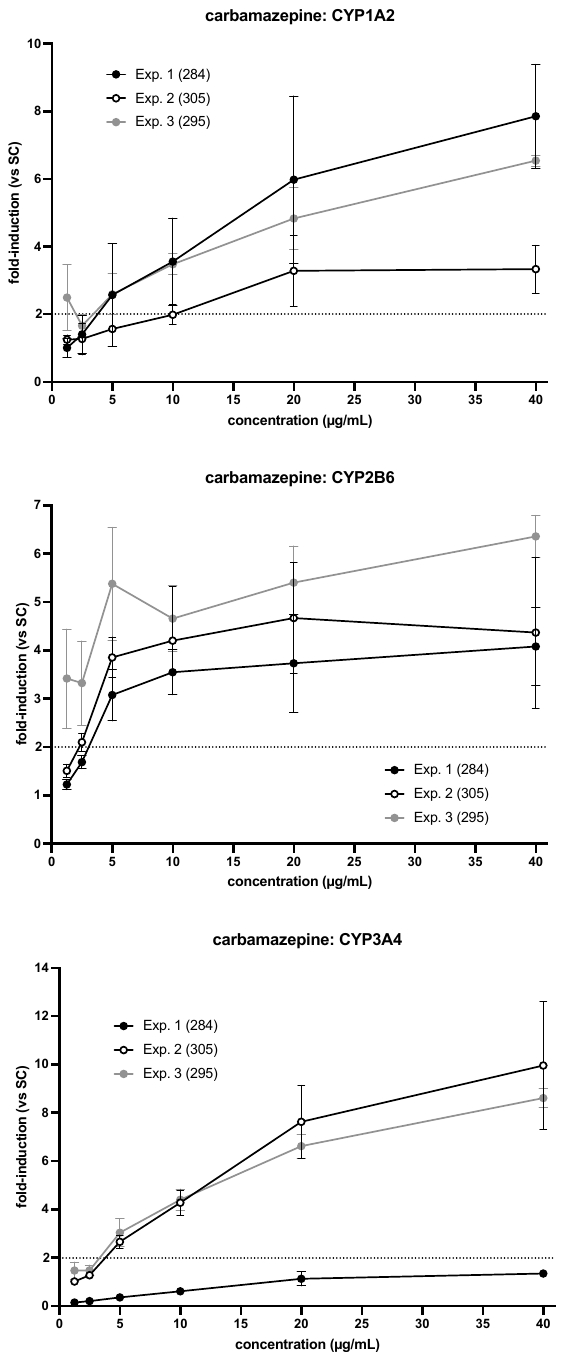

**Phenytoin**

**INRAE	UU**

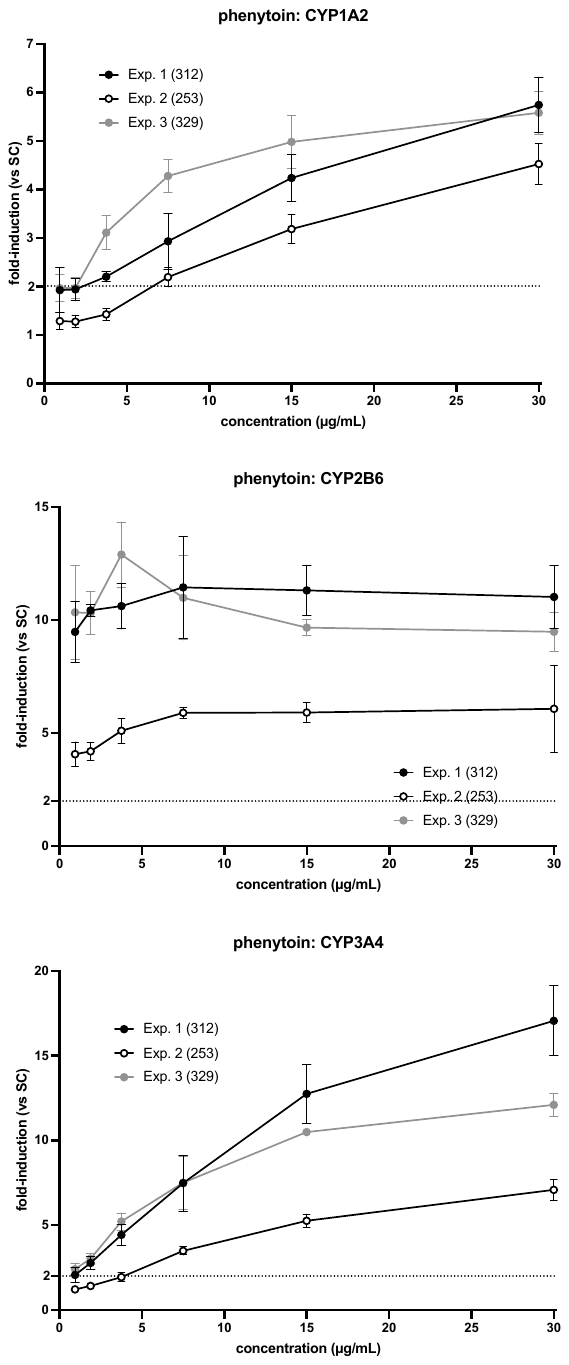

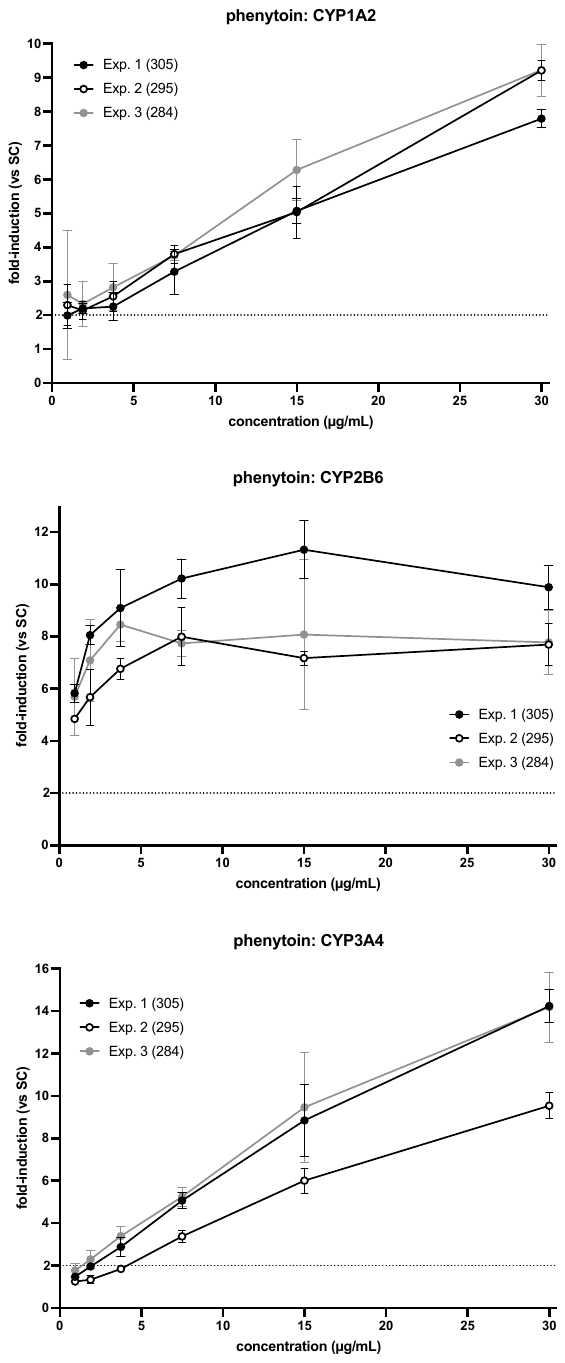

**Penicillin G**

**INRAE	UU**

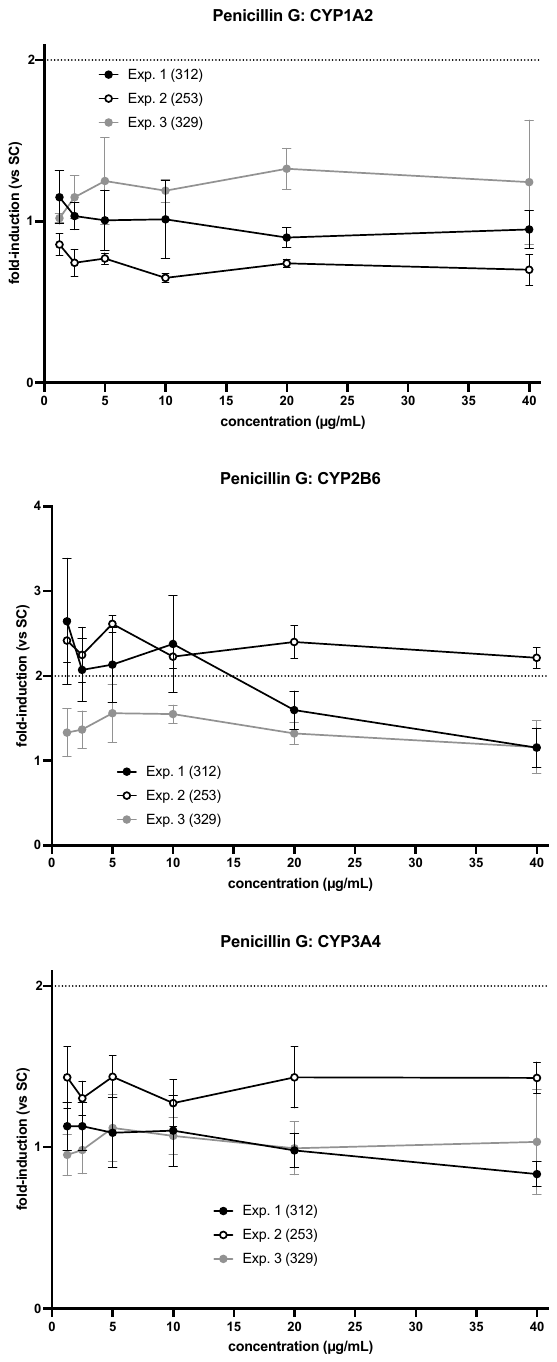

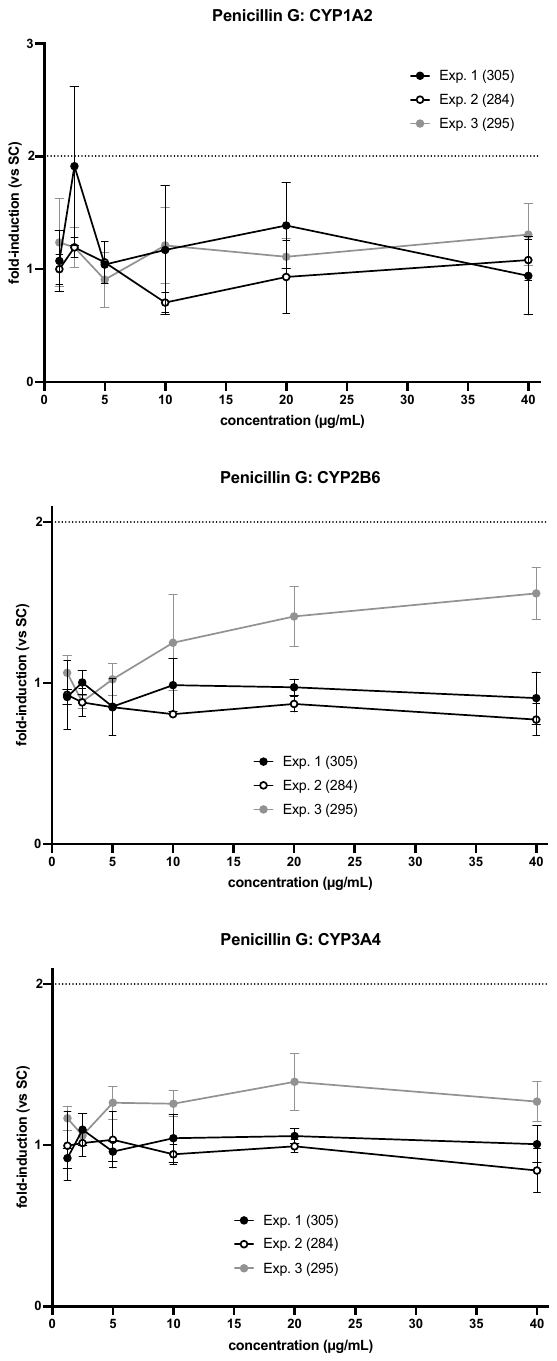

**Sulfinpyrazone**

**INRAE	UU**

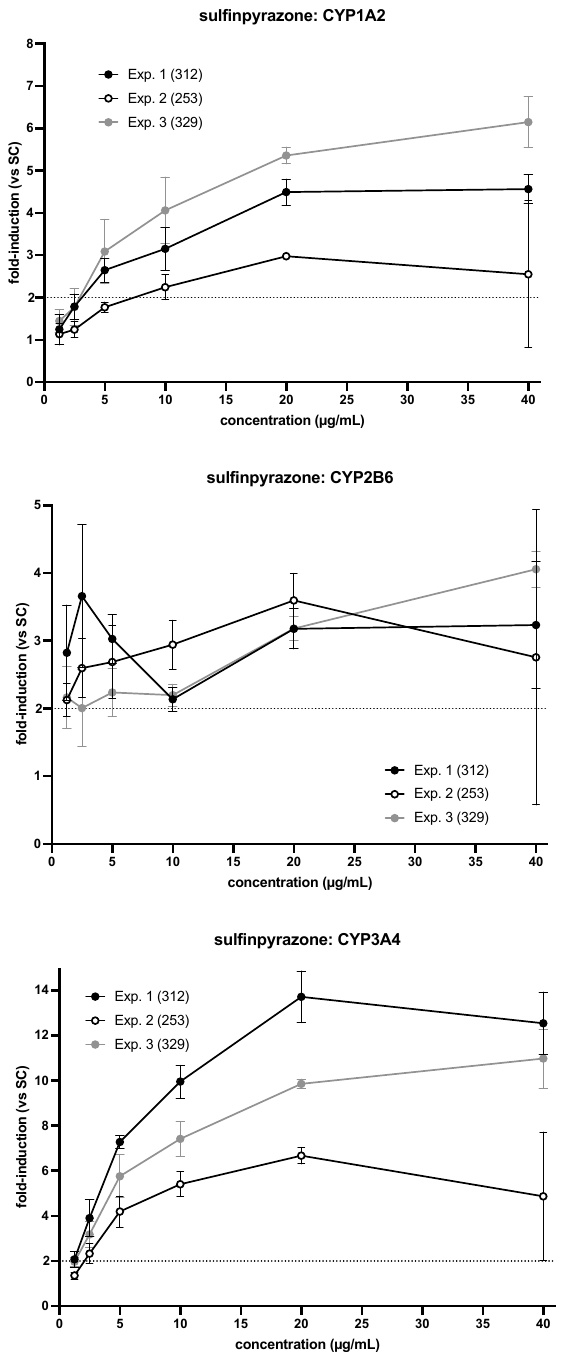

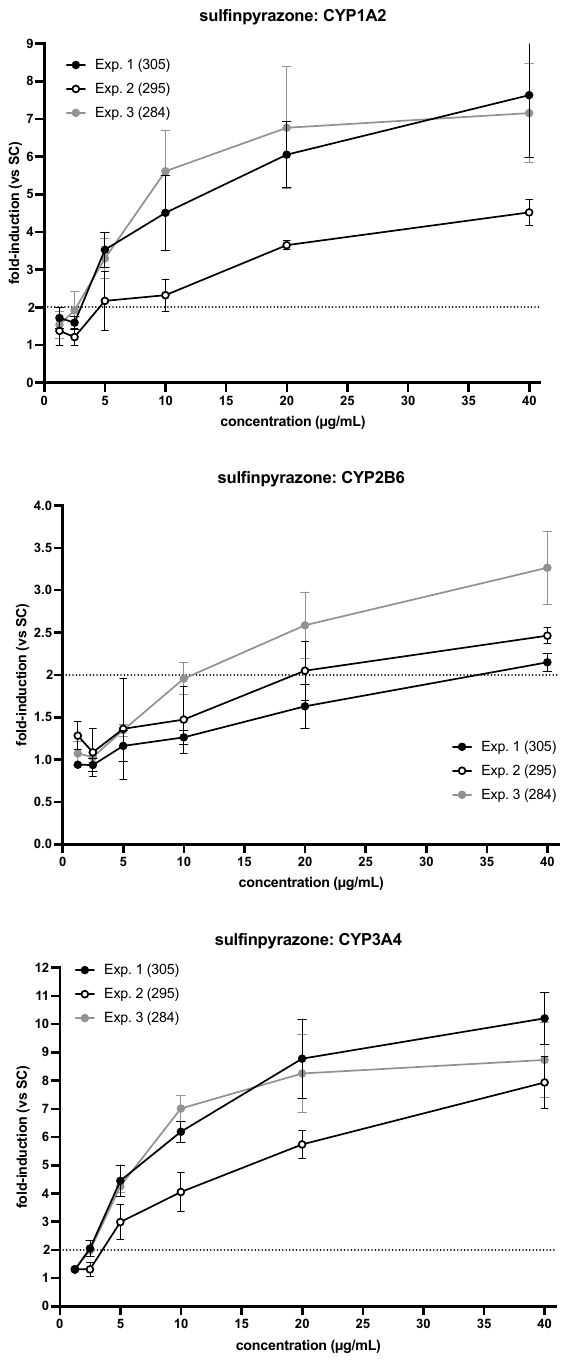

**Bosentan**

**INRAE	UU**

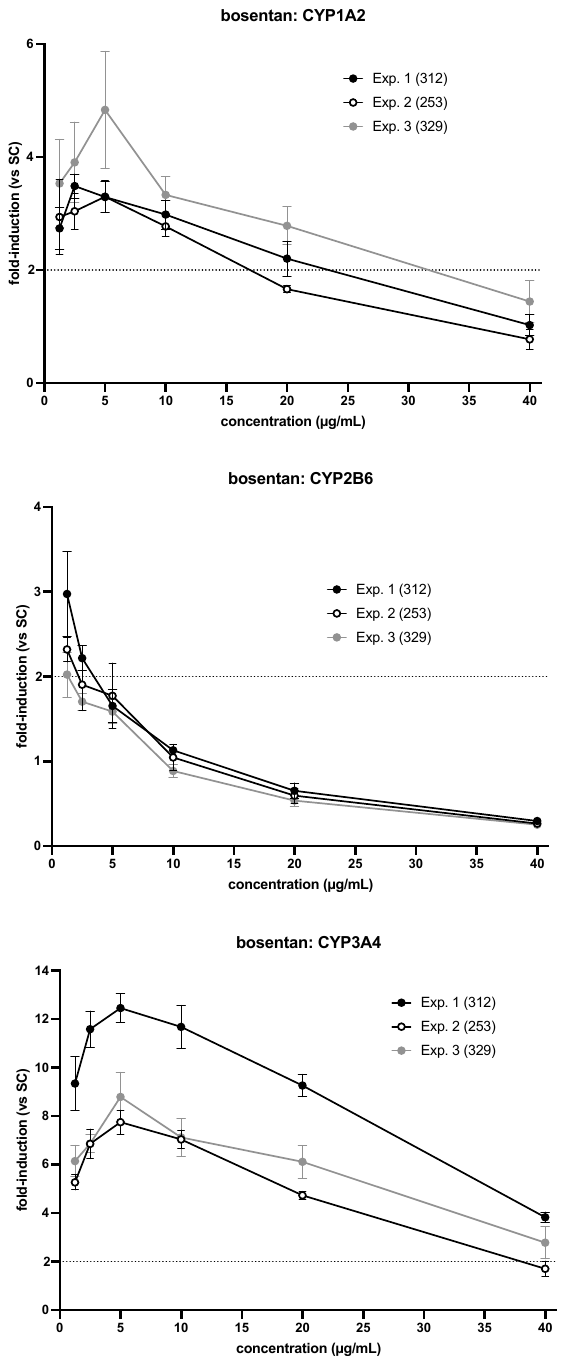

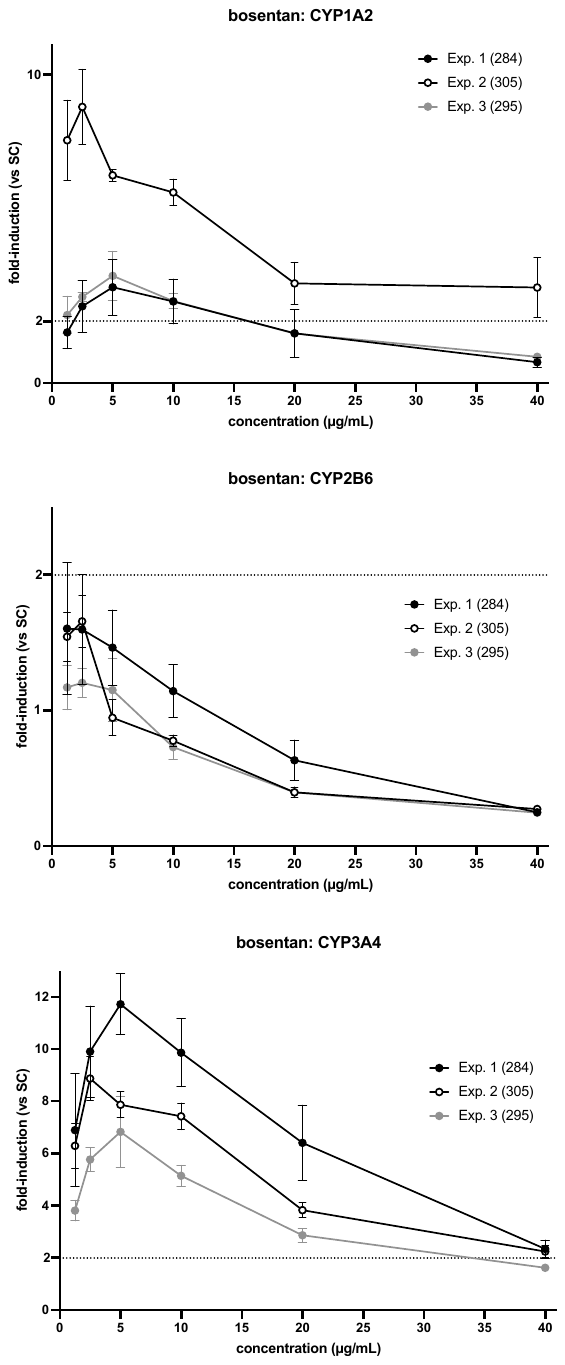

**Artemisinin**

**INRAE	UU**

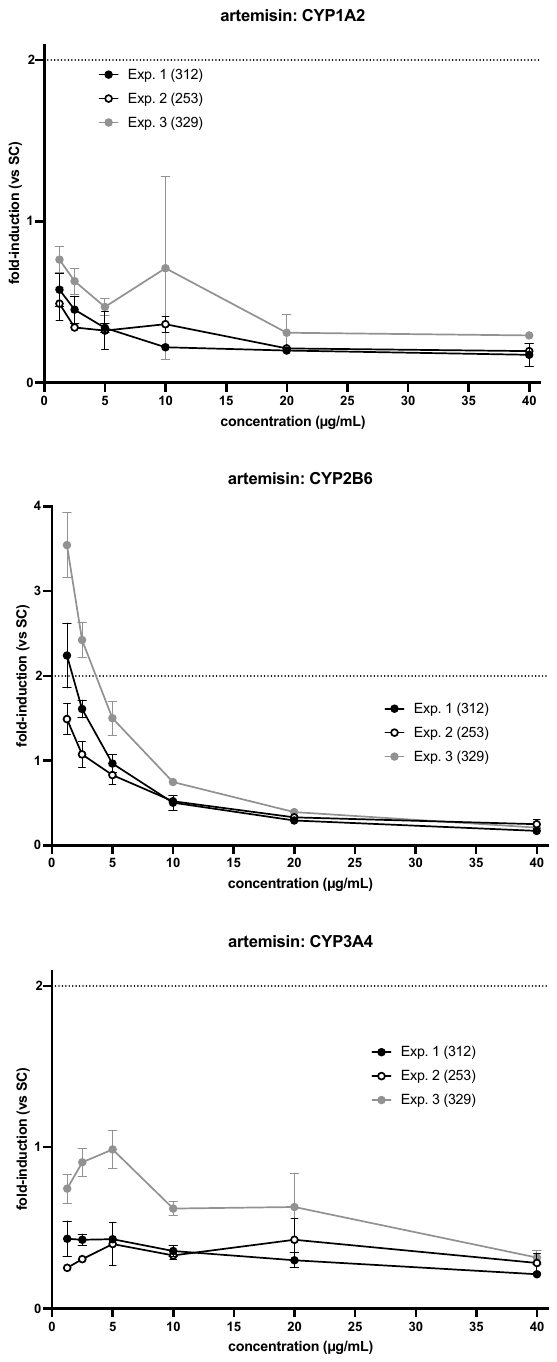

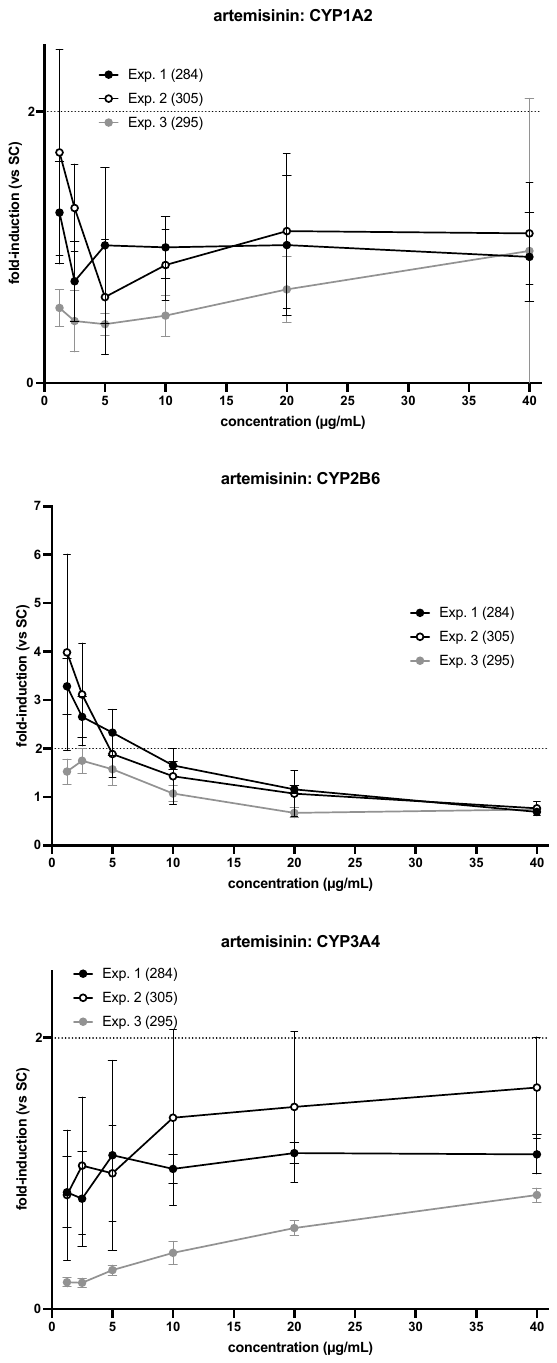

**Rifampicin**

INRAE	UU

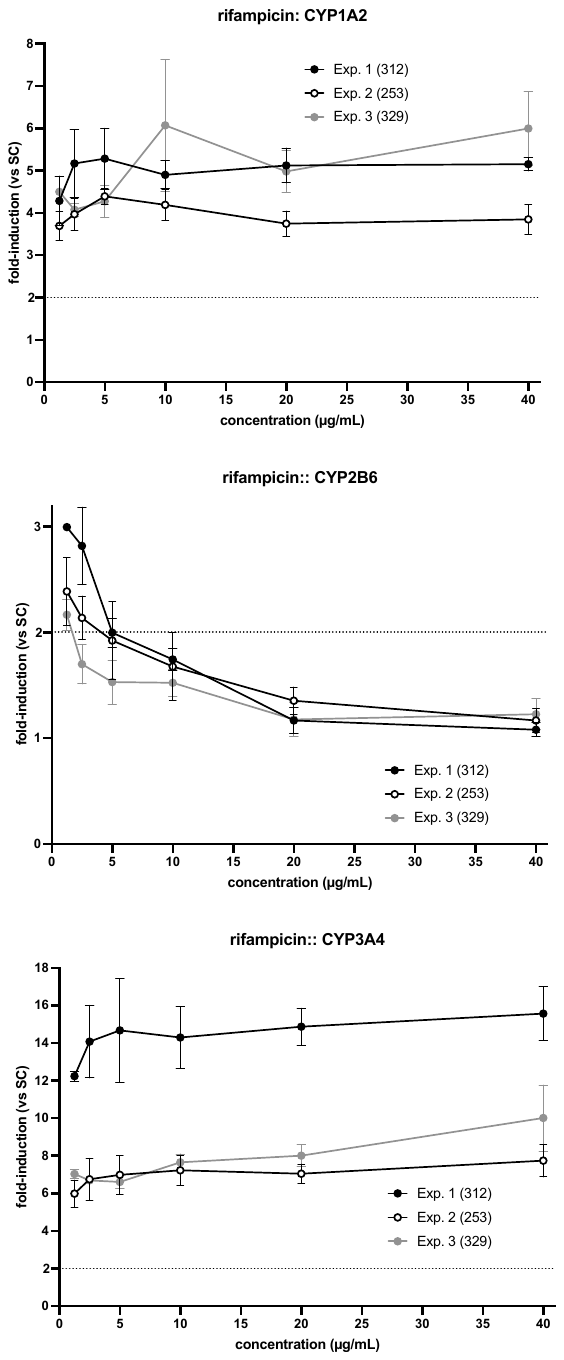

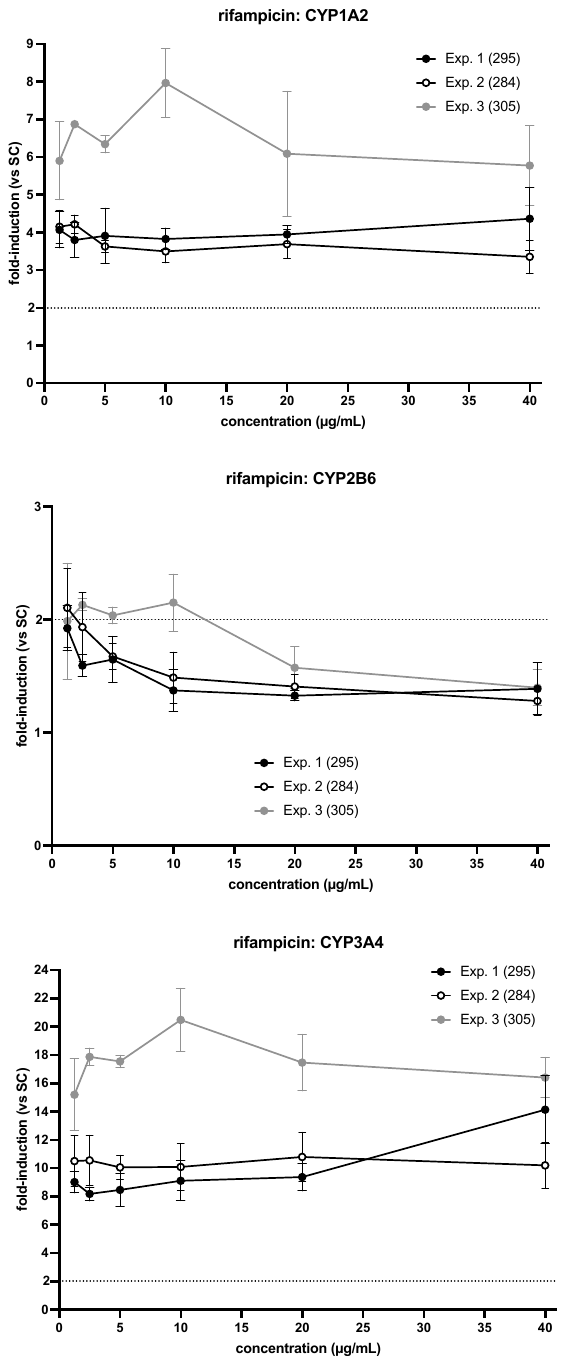

**Metropolol**

**INRAE	UU**

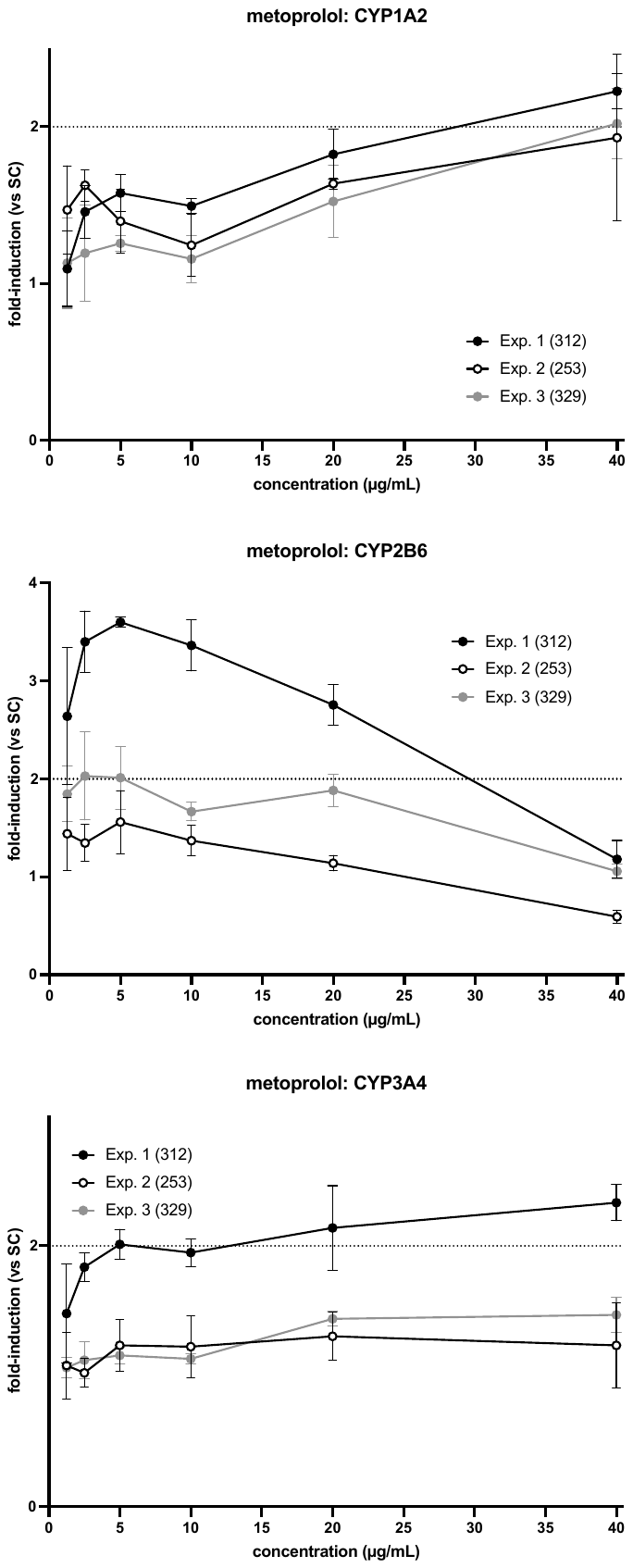

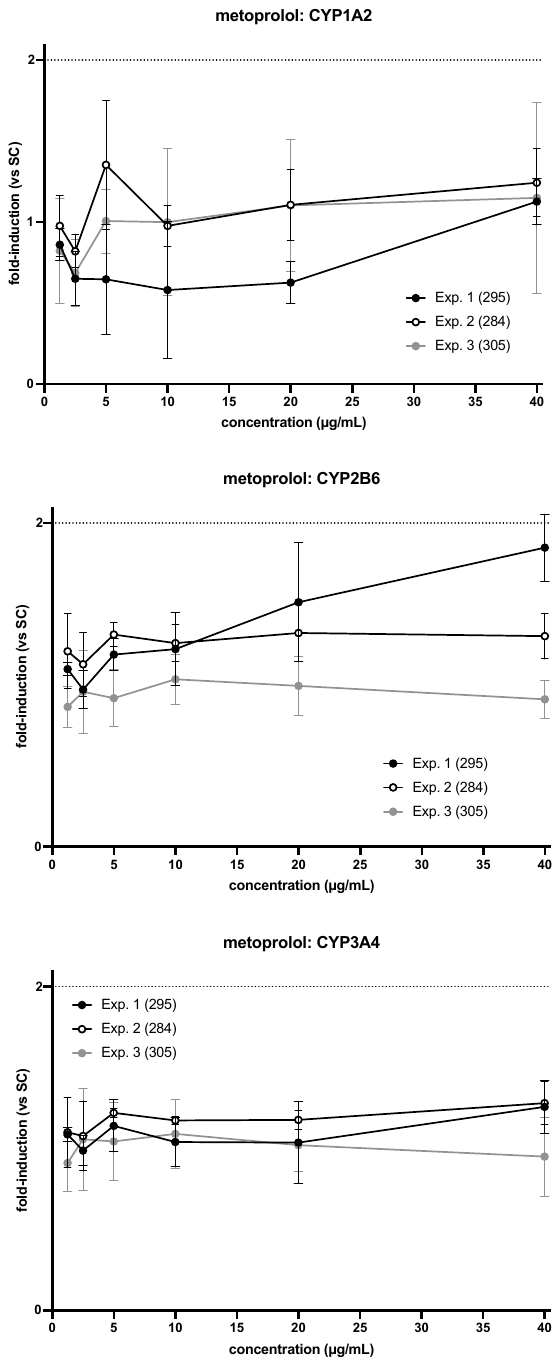

**Sotalol**

**INRAE	UU**

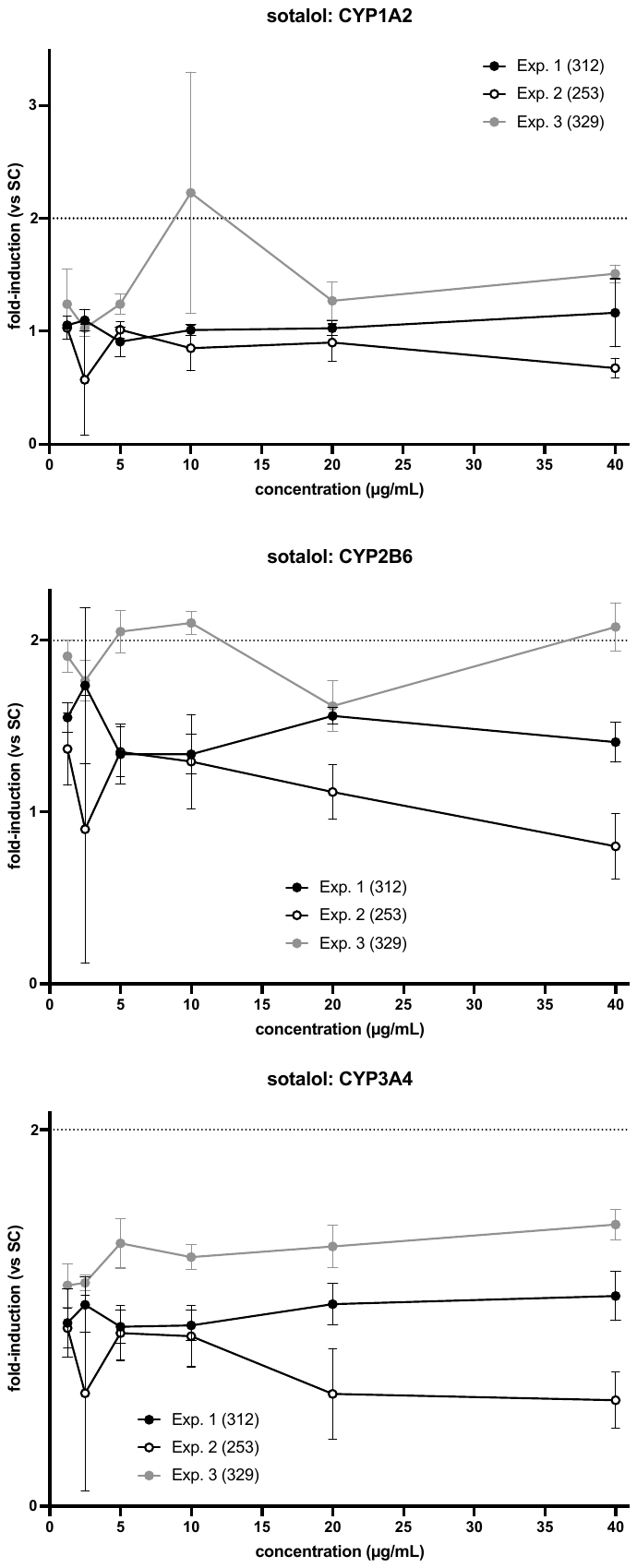

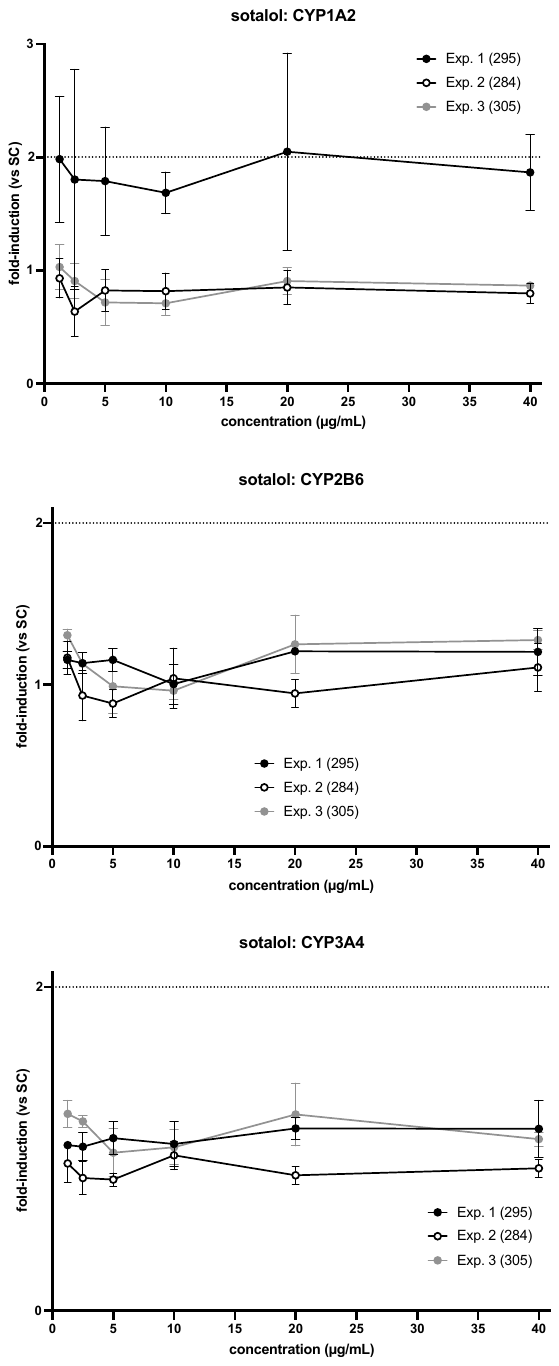

**Supplementary Information 6**

**Graphical representation of the results for the 6 augmentation chemicals**

**Permethrin**

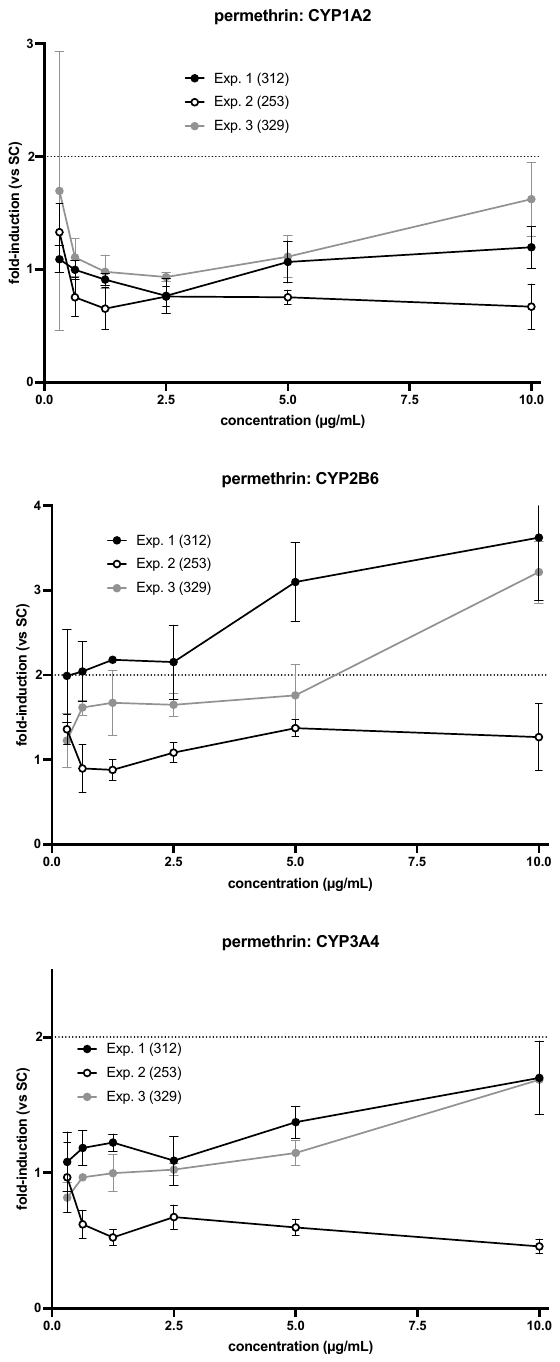
**INRAE	UU**

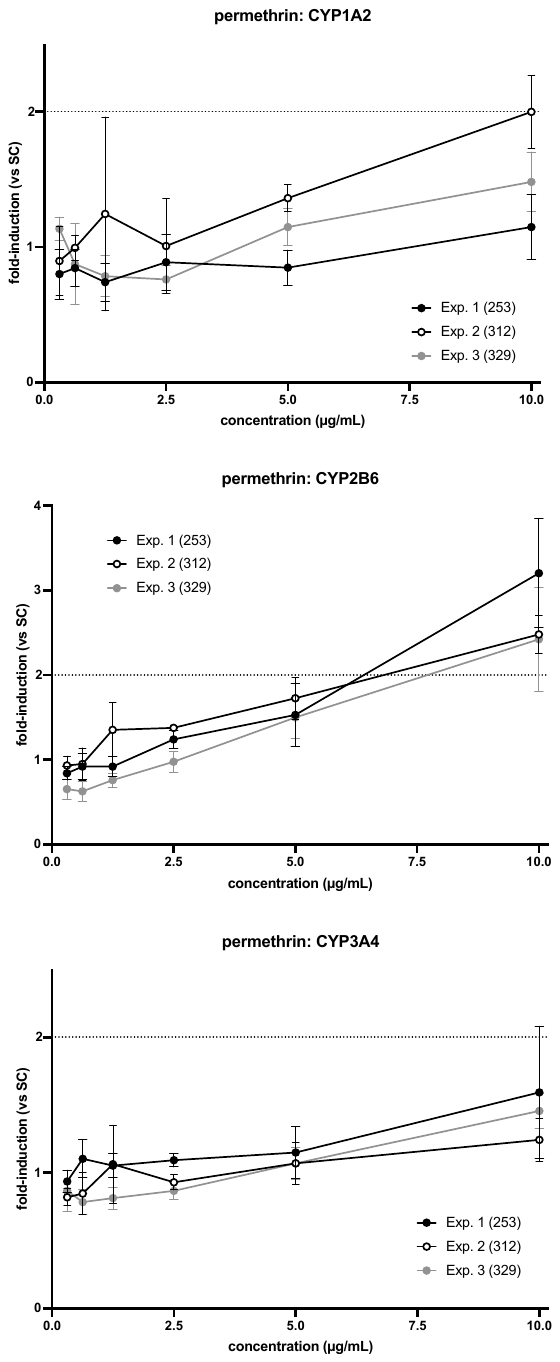

**Fipronil**

**INRAE	UU**

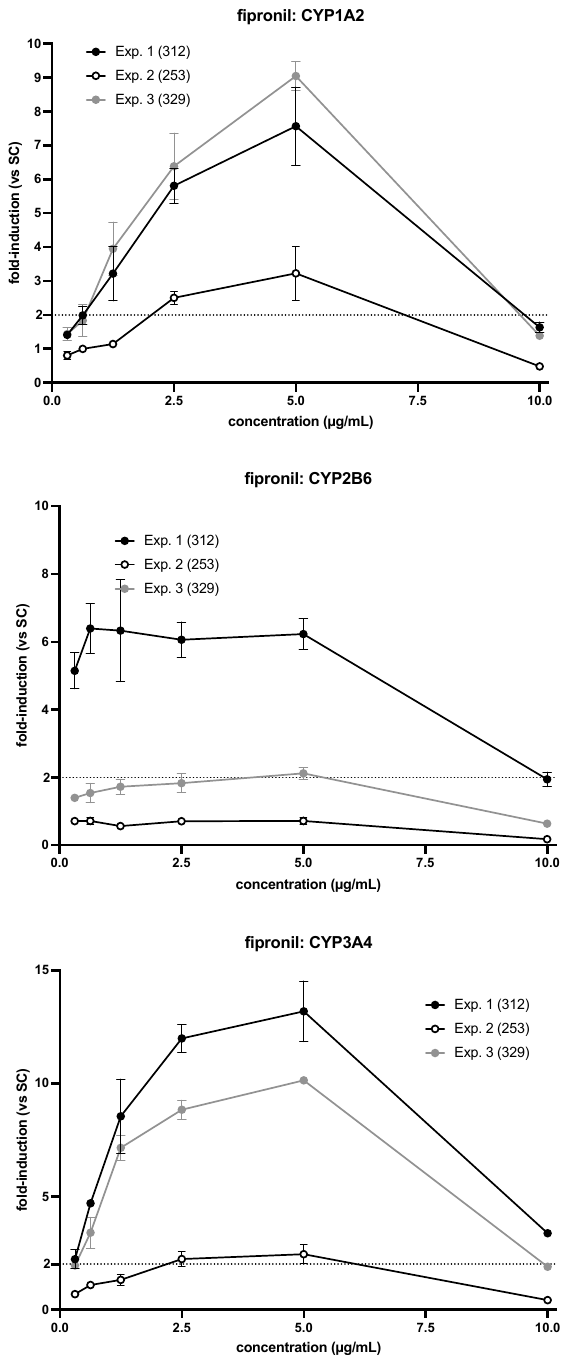

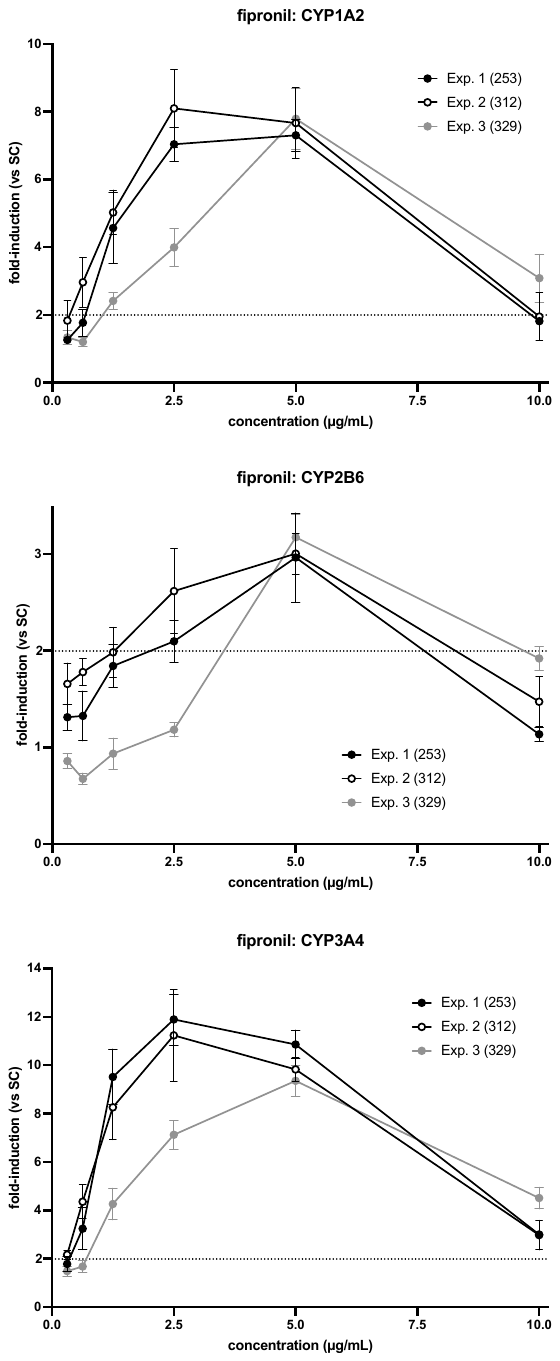

**Tebuconazole**

**INRAE	UU**

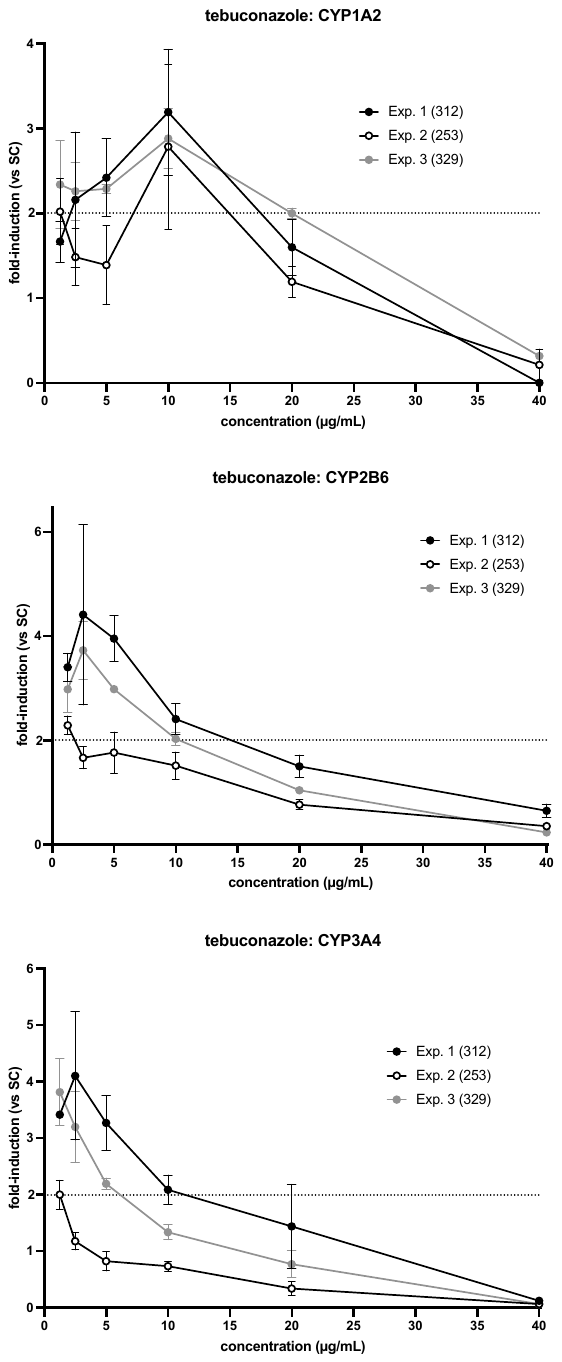

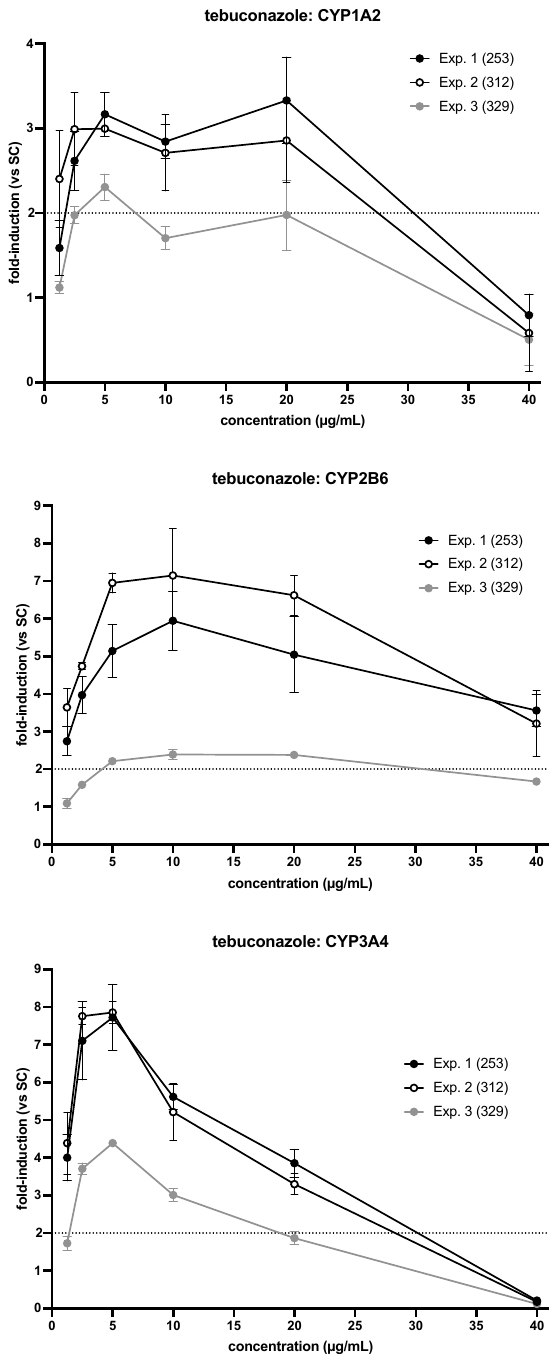

**DEET**

INRAE	UU

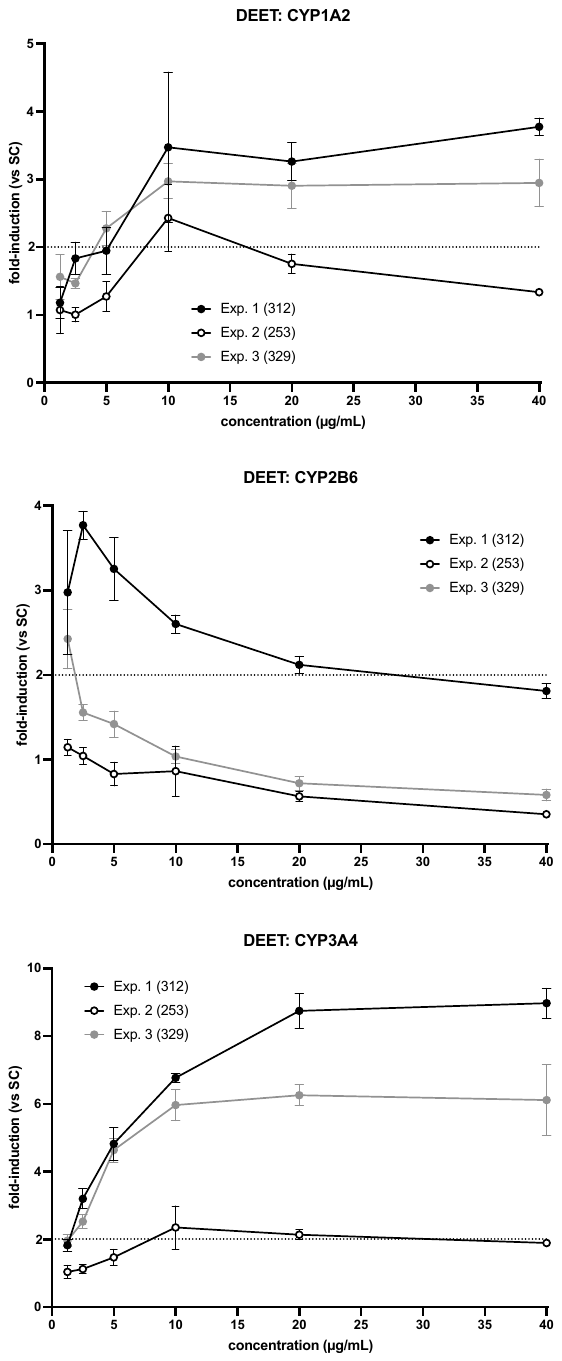

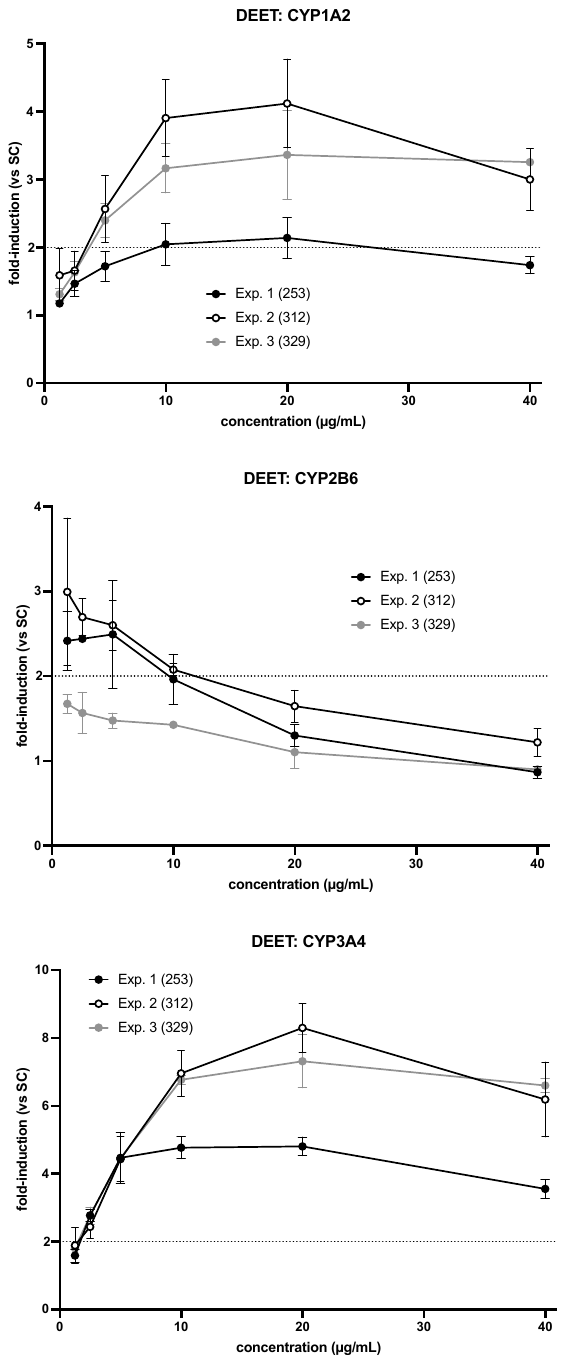

**Chlorpyrifos**

**INRAE	UU**

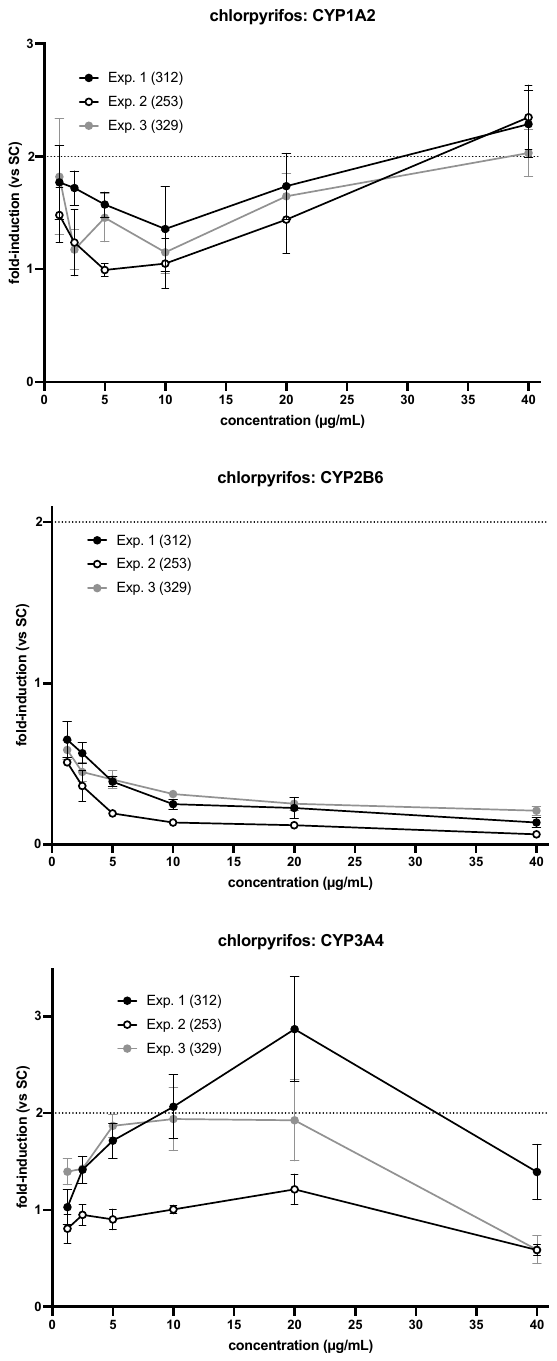

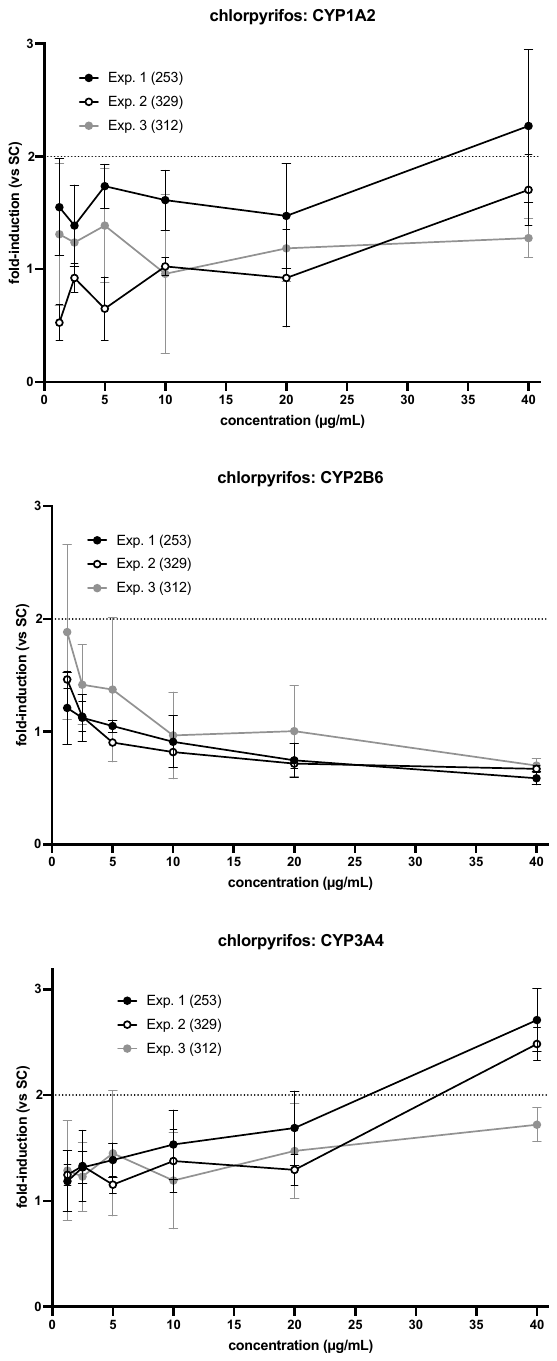

**Benfuracarb**

**INRAE	UU**

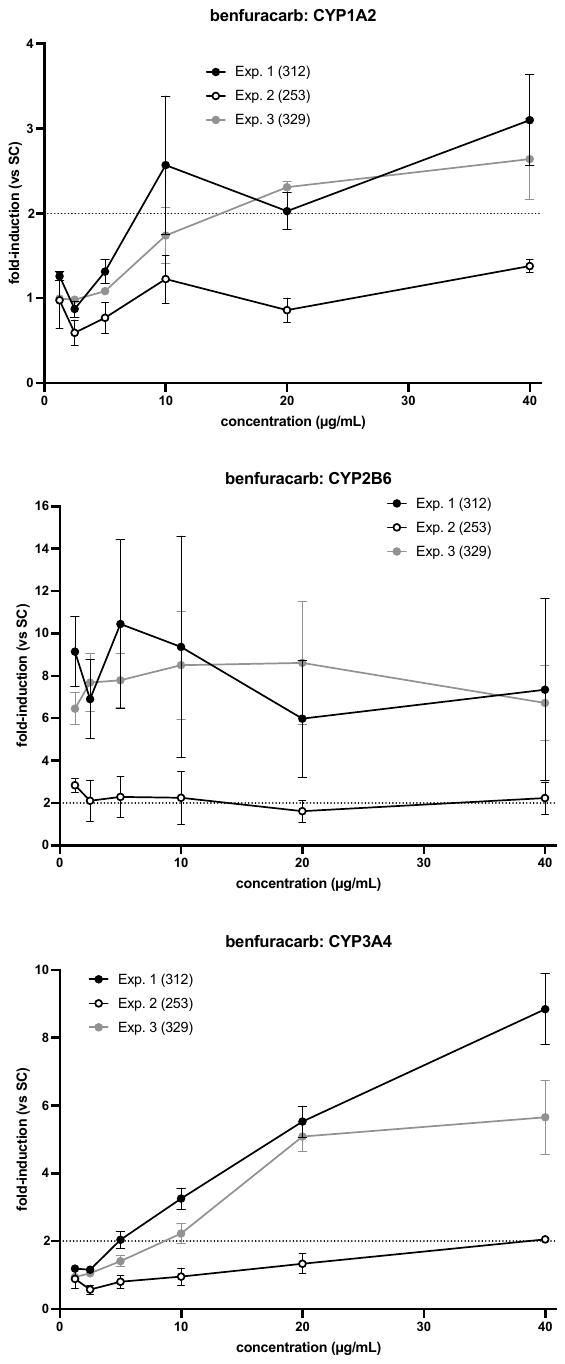

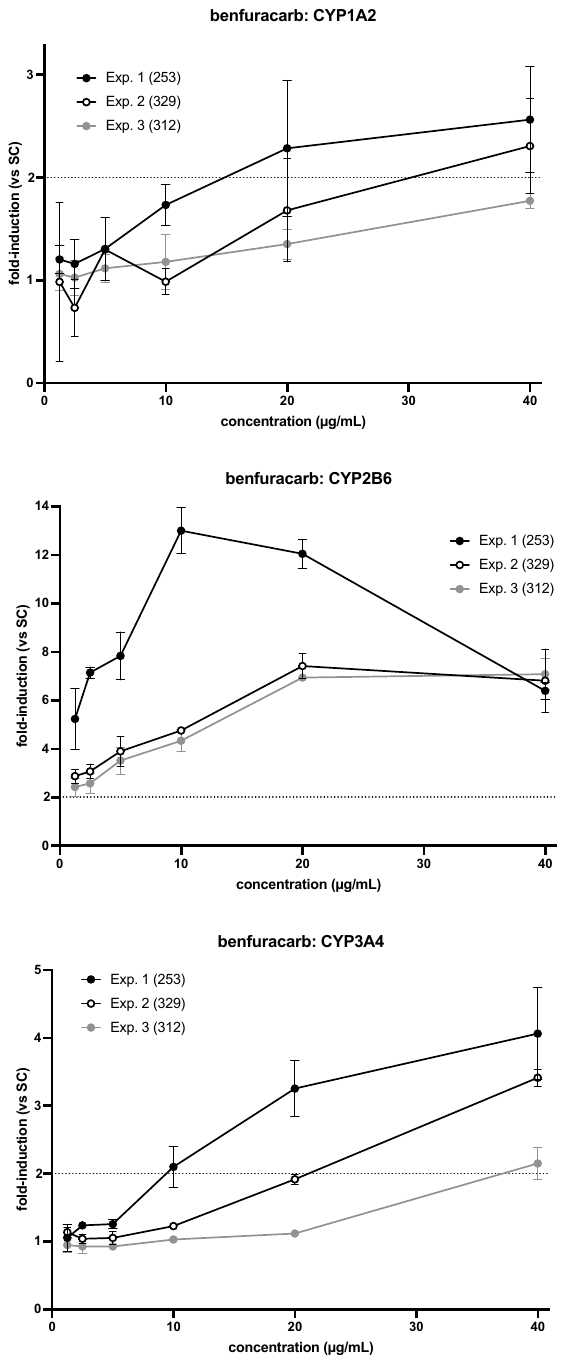
